# Complex genetic architecture of three-dimensional craniofacial shape variation in domestic pigeons

**DOI:** 10.1101/2021.03.15.435516

**Authors:** Elena F. Boer, Emily T. Maclary, Michael D. Shapiro

## Abstract

Deciphering the genetic basis of vertebrate craniofacial variation is a longstanding biological problem with broad implications in evolution, development, and human pathology. One of the most stunning examples of craniofacial diversification is the adaptive radiation of birds, in which the beak serves essential roles in virtually every aspect of their life histories. The domestic pigeon (*Columba livia*) provides an exceptional opportunity to study the genetic underpinnings of craniofacial variation because of its unique balance of experimental accessibility and extraordinary phenotypic diversity within a single species. We used traditional and geometric morphometrics to quantify craniofacial variation in an F_2_ laboratory cross derived from the straight-beaked Pomeranian Pouter and curved-beak Scandaroon pigeon breeds. Using a combination of genome-wide quantitative trait locus scans and multi-locus modeling, we identified a set of genetic loci associated with complex shape variation in the craniofacial skeleton, including beak curvature, braincase shape, and mandible shape. Some of these loci control coordinated changes between different structures, while others explain variation in the size and shape of specific skull and jaw regions. We find that in domestic pigeons, a complex blend of both independent and coupled genetic effects underlie three-dimensional craniofacial morphology.

## Introduction

The vertebrate skull serves essential roles in numerous biological processes, including respiration, feeding, communication, and protecting the brain and sense organs. Throughout vertebrate evolution, dramatic diversification of craniofacial morphology has accompanied successful occupation of diverse ecological and dietary niches. Identifying the genetic programs that underlie variation in the form and function of the craniofacial complex is a longstanding goal with implications in diverse biological fields, including evolutionary biology, ecology, embryology, molecular biology, and genetics. In addition, deciphering the genetic basis of craniofacial variation represents an important clinical objective, as many human craniofacial disorders are caused by genetic mutations that disrupt morphogenesis and result in phenotypes that fall outside of the spectrum of normal variation (Trainor 2010; Twigg and Wilkie 2015).

Studies of the genetic basis of vertebrate craniofacial variation often focus on traits with a relatively simple genetic basis and/or represent complex craniofacial variation as simplified measurements. For example, in wild species of birds, researchers have identified genes that are putatively associated with simple measures of beak variation, such as overall size (*IGF1*) in Black-bellied seedcrackers (vonHoldt *et al*. 2018); length (*COL4A5*) in great tits (Bosse *et al*. 2017); and length (*CALM1*), width (*BMP4*), and overall size (*ALX1*, *HMGA2*) in Darwin’s finches (Abzhanov 2004; Abzhanov *et al*. 2006; Mallarino *et al*. 2011; Lamichhaney *et al*. 2015, 2016). Our understanding of the genetic architecture of 3D craniofacial shape remains comparatively limited, in part because of the inherent challenges of quantifying complex morphological variation and implementing forward genetic approaches to map the underlying genetic architecture. A number of recent studies use 3D phenotypes and genetic mapping to determine the architecture of craniofacial variation in several vertebrates, including dogs, cichlids, mice, and humans (Albertson *et al*. 2003, 2005b; Roberts *et al*. 2011; Schoenebeck *et al*. 2012; Powder *et al*. 2014; Pallares *et al*. 2015; Shaffer *et al*. 2016; Marchant *et al*. 2017; Claes *et al*. 2018; Xiong *et al*. 2019; Katz *et al*. 2020). A consistent take-home message from this body of work is that the craniofacial skeleton and its underlying genetic architecture is remarkably complex; in many cases, multiple genetic loci explain only a small percentage of overall craniofacial shape variation. Sometimes, the major genetic or developmental controls of variation appear to be unique to a particular species or population, while others show overlap among species (e.g., BMP signaling in birds, cichlids, and dogs (Abzhanov 2004; Albertson *et al*. 2005a; Schoenebeck *et al*. 2012)).

The massive diversity of craniofacial morphology among birds has inspired excellent comparative morphometric analyses of shape variation across species (recent examples include (Campàs *et al*. 2010; Mallarino *et al*. 2012; Fritz *et al*. 2014; Bright *et al*. 2016, 2019; Cooney *et al*. 2017; Young *et al*. 2017; Felice and Goswami 2018; Yamasaki *et al*. 2018; Navalón *et al*. 2019, 2020)). In contrast, there are few examples of pairing geometric morphometric shape analysis with genome-wide scans to identify the genetic architecture of avian craniofacial variation (but see (Yusuf *et al*. 2020)). The domestic pigeon (*Columba livia*) provides an extraordinary opportunity to disentangle the genetic architecture of complex craniofacial variation. Pigeons have spectacular craniofacial variation among hundreds of breeds within a single species; the magnitude of their intraspecific diversity is more typical of interspecific diversity (Baptista *et al*. 2009). Recently, Young et al. (Young *et al*. 2017) used geometric morphometrics to compare craniofacial shape among breeds of domestic pigeon and diverse wild bird species and concluded that the shape changes that differentiate pigeon breeds recapitulate the major axes of variation in distantly related wild bird species. However, unlike most distantly related species, domestic pigeon breeds are interfertile, so we can establish laboratory crosses between anatomically divergent forms and map the genetic architecture of variable traits.

The goal of this study is to identify the genetic architecture of craniofacial shape variation in an F_2_ population derived from pigeon breeds with dramatically different craniofacial morphologies. First, we report traditional linear measurements that define the height, width, and depth of three craniofacial substructures: the upper beak, braincase, and mandible. Then, we use geometric morphometrics to quantify three-dimensional shape variation in these three substructures. Finally, we use these morphological data to perform genome-wide QTL scans and multi-locus modeling to map the genetic architecture of complex craniofacial variation, including beak curvature.

## Results

To identify the genetic architecture underlying craniofacial shape variation in domestic pigeons, we performed an F_2_ intercross between a male Pomeranian Pouter (Pom) and two female Scandaroons (Scan) (Figure 1A-D, Supplemental Figure 1). These two breeds display highly divergent craniofacial morphologies, in addition to other variable phenotypes (e.g., plumage color, hindlimb epidermal appendages (Domyan *et al*. 2014, 2016)). The Pom breed has a straight beak that is qualitatively similar to the beak of many other domestic pigeon breeds, as well as the ancestral rock pigeon (Figure 1A,C, Supplemental Figure 1). In contrast, the curved beak of the Scandaroon breed is one of the most extreme craniofacial phenotypes observed in any domestic pigeon breed (Figure 1B,D, Supplemental Figure 1).

**Figure 1.**
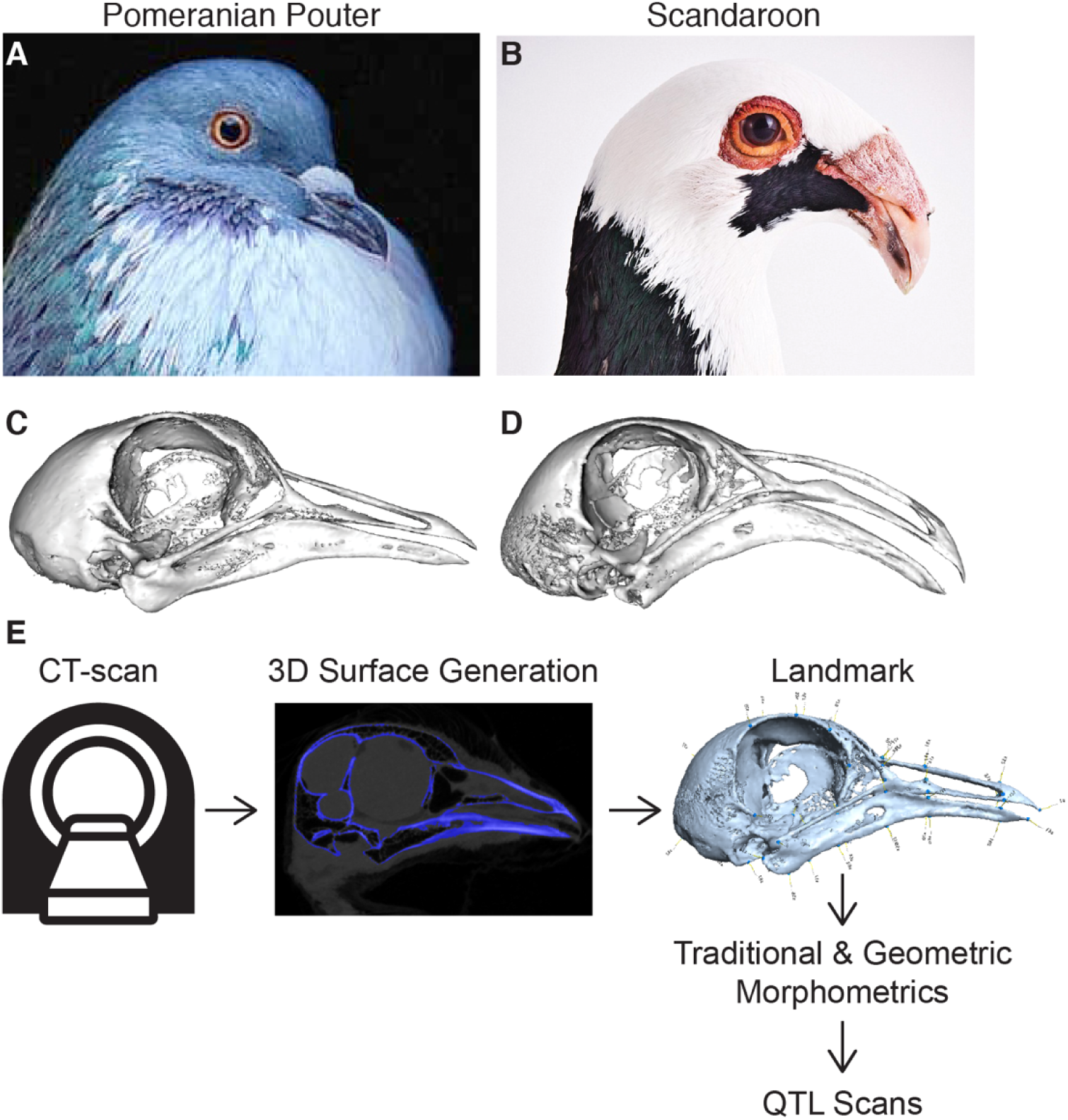
Morphometric analyses of craniofacial shape and quantitative trait loci (QTL) mapping in a pigeon F_2_ intercross. (A-B) Representative images of the Pomeranian Pouter (Pom, A) and Scandaroon (Scan, B) breeds of domestic pigeon used to generate the Pom × Scan F2 intercross. (C-D) 3D surface models of the craniofacial skeletons of the male Pom (C) and one of the female Scan (D) cross founders. (E) Experimental approach to identify genetic architecture of craniofacial variation in the Pom × Scan cross. Image credits (used with permission): Drew Snyder (A); Richard Bailey (B).

To visualize and quantify variation in the Pom × Scan F_2_ population, we scanned the cross founders and 116 F_2_ individuals using micro-computed tomography (micro-CT) and generated three-dimensional surface models of the craniofacial skeleton (Figure 1E). We developed an atlas of 73 landmarks that collectively define the shape of the upper beak, braincase, and mandible (Supplemental Figure 2, Supplemental Table 1) and applied the landmark set to the cross founders and all F_2_ individuals.

### Morphometric analyses of linear dimensions

We first measured 10 linear distances between landmark pairs that define the length, width, and depth of three skull and jaw substructures – upper beak, braincase, and mandible – to quantify variation in the Pom × Scan F_2_ population (Supplemental Table 2). We found that all linear measurements are normally distributed within the population, with the exception of rostral mandible width (Supplemental Figure 3). To determine if craniofacial size and shape are predicted by body size, we performed a linear regression of each linear measurement on body mass, a commonly-used proxy for body size ((Hallgrímsson *et al*. 2019); Supplemental Figure 4). Most (8/10) skull and jaw linear measurements had a significant and positive allometric association with body size; only braincase length and width were independent of body size (Supplemental Figure 4). After extracting non-allometric variation, we compared the residuals of each linear measurement between sexes and found that males had significantly longer and deeper craniofacial structures relative to females (Supplemental Figure 4). In contrast, among the measurements of craniofacial width, only rostral braincase and caudal mandible width were sex-dependent (Supplemental Figure 4). These results demonstrate that both allometric and non-allometric shape variation exist within the Pom × Scan F_2_ population, and that craniofacial length and depth are regulated in part by a sex-linked factor that has only a limited effect on width.

### QTL on 5 linkage groups are associated with linear variation in craniofacial structures

To identify genomic regions associated with variation in craniofacial length, width, and depth, we performed genome-wide quantitative trait locus (QTL) scans for each of the 10 linear measurements. We identified significant major-effect QTL for 6 linear measurements representing all three skull and jaw substructures (Table 1), including upper beak width and depth (Figure 2), braincase length and width (Supplemental Figure 5), and mandible length and width (Supplemental Figure 6). Two of the major-effect QTL (LG1 and LG8) are especially notable because they control variation in correlated traits.

**Figure 2.**
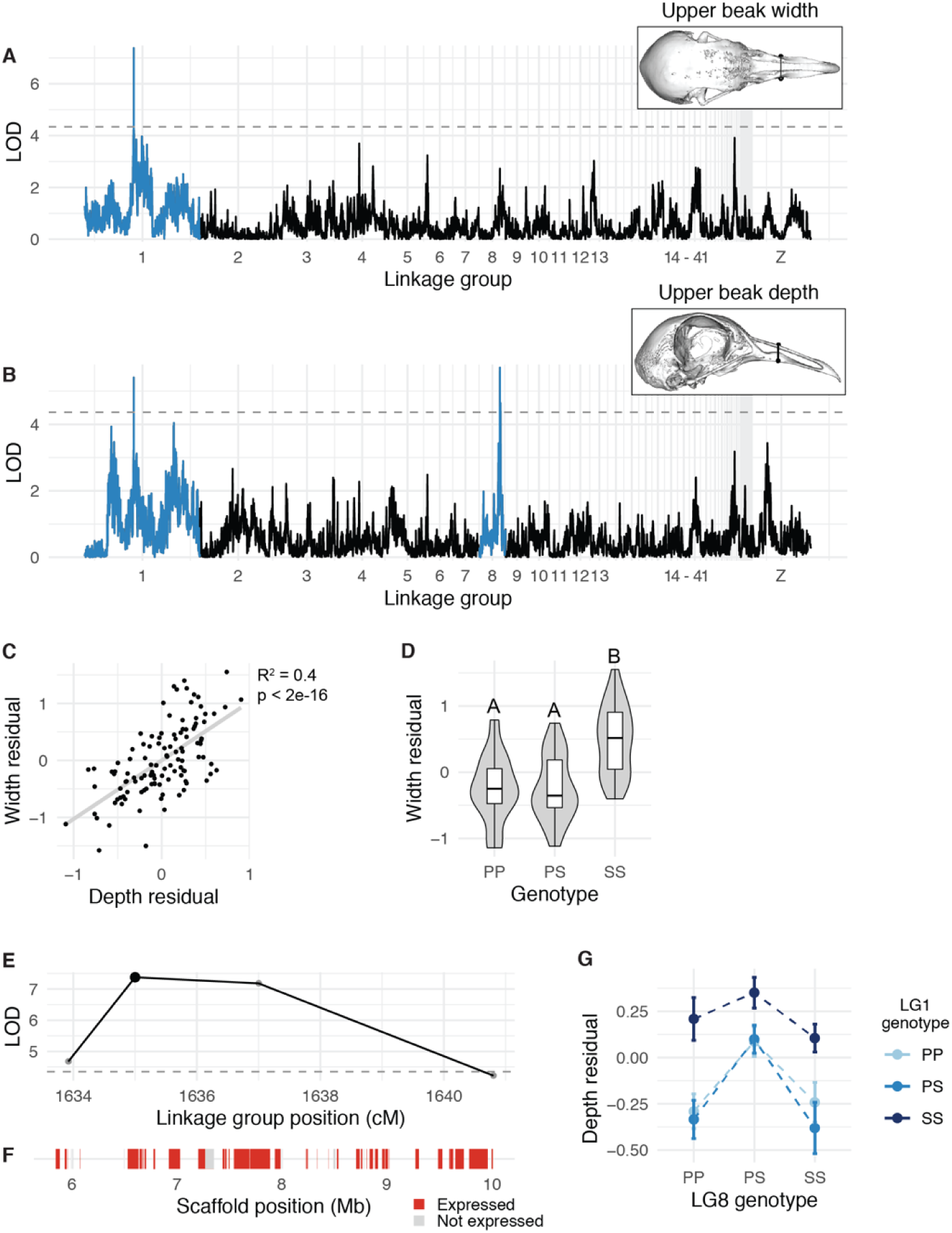
QTL associated with upper beak width and depth. (A-B) Genome-wide QTL scans for upper beak width (A) and depth (B). Dashed horizontal line indicates 5% genome-wide significance threshold and linkage groups with significant QTL peaks are highlighted in blue. (C) Scatterplot of upper beak width and depth measurements for all Pom × Scan F_2_ individuals. Plotted values are residuals from regression on body mass. (D) Beak width effect plot. Letters denote significance groups, p-values determined via Tukey test: PP vs. SS = 4.3e-06, PS vs. SS = 9.1e-06. (E) LOD support interval for beak width QTL scan. Dots indicate linkage map markers; the larger black dot highlights the peak marker that was used to estimate QTL effects in (D). (F) Genes located within LOD support interval, color coded based on expression status in HH29 facial primordia. (G) Interaction plot between LG1 and LG8 QTL associated with upper beak depth. P = allele from Pom founder, S = allele from Scan founder.

**Table 1.** QTL associated with skull and jaw linear measurements and shape.

| QTL | LG | Position<br>(cM) | LOD | PVE<br>(%) | Interval size<br>(Mb) | Total<br>genes | Expressed<br>genes |
| --- | --- | --- | --- | --- | --- | --- | --- |
| *Linear measurements* |  |  |  |  |  |  |  |
| Upper beak width | 1 | 1635.00 | 7.38 | 25.39 | 4.21 | 41 | 33 |
| Upper beak depth | 1 | 1635.00 | 5.41 | 19.32 | 4.21 | 41 | 33 |
| Upper beak depth | 8 | 688.81 | 5.71 | 20.27 | 0.32 | 5 | 5 |
| Braincase length | 2 | 1082.73 | 5.56 | 19.81 | 50.89 | 446 | 399 |
| Braincase width<br>(caudal) | 5 | 680.84 | 4.65 | 16.86 | 0.48 | 5 | 5 |
| Mandible length | 10 | 236.20 | 5.05 | 18.16 | 0.88 | 26 | 24 |
| Mandible width | 8 | 699.06 | 6.41 | 22.46 | 0.09 | 2 | 2 |
| *Shape* |  |  |  |  |  |  |  |
| UBB PC2 | 3 | 1361.00 | 4.93 | 17.77 | 17.34 | 171 | 146 |
| UBB PC3 | 13 | 454.00 | 4.53 | 16.45 | 1.30 | 4 | 3 |
| UBB PC4 | 10 | 614.97 | 4.78 | 17.29 | 10.19 | 52 | 45 |
| UBB PC4 | 11 | 426.06 | 4.57 | 16.59 | 15.99 | 209 | 177 |
| MAN PC3 | 2 | 716.24 | 4.58 | 16.62 | 1.94 | 27 | 21 |
| MAN PC3 | 3 | 1432.70 | 5.75 | 20.41 | 7.20 | 35 | 31 |
| MAN PC4 | 11 | 15.00 | 6.42 | 22.51 | 1.42 | 34 | 21 |
| MAN PC5 | 20 | 391.81 | 5.01 | 18.05 | 0.54 | 6 | 6 |

#### A QTL on LG1 is associated with beak width and depth

Upper beak width and depth are significantly positively associated in the cross (R^2^ = 0.4, p < 2e-16, Figure 2C). Perhaps not surprisingly, both measurements mapped to the same QTL on LG1 (upper beak width: LOD = 7.4, PVE = 25.4%, Figure 2A; upper beak depth: LOD = 5.4, PVE = 19.3%, Figure 2B). The LG1 Pom allele is dominant, as upper beak width and depth of heterozygotes are indistinguishable from Pom homozygotes (Figure 2D). F_2_ individuals homozygous for the Scan allele had significantly wider and deeper upper beaks than individuals homozygous for the Pom allele (Figure 2D).

The LG1 LOD support interval is a 4.16-Mb region that includes 41 protein-coding genes (Figure 2E-F). To prioritize candidate genes within the interval, we cross-referenced the gene list to RNA expression data from pigeon facial primordia from the Racing Homer breed (developmental stage equivalent to Hamburger-Hamilton chicken stage 29, or HH29; (Hamburger and Hamilton 1951)). Of the 41 genes in the upper beak width/depth interval, 33 genes are expressed in the developing pigeon face (Figure 2F, Supplemental Table 3). Notably, *FGF6* is located near the center of the QTL interval (34 kb downstream of the LG1 peak marker). *FGF6* is expressed in craniofacial structures during chicken embryogenesis (Kumar and Chapman 2012), and *Fgf6^-/-^* mutant mice have shorter snouts than their wildtype littermates (Floss *et al*. 1997), demonstrating a role for this gene in outgrowth of vertebrate facial structures.

#### A QTL on LG8 is associated with beak depth and mandible width

A second major-effect QTL on LG8 was associated with upper beak depth (LOD = 5.7, PVE = 20.3%), but not width (Figure 2B). F_2_ heterozygotes have a wider beak than either homozygote (Figure 2G). The LG8 QTL functions additively with the LG1 QTL described above: two copies of the LG1 Scan allele increased beak width for all LG8 genotypes (Figure 2G). The 0.36-Mb LOD support interval on LG8 contains only 5 genes (*USP33*, *ZZZ3*, *AK5*, *PIGK*, *ST6*), all of which are expressed in embryonic pigeon craniofacial tissues (Supplemental Figure 7, Supplemental Table 4), but none are known to play a role in craniofacial development in other species.

A major-effect QTL associated with mandible width overlaps with the upper beak depth QTL on LG8 (LOD = 6.4, PVE = 22.5%, Supplemental Figure 6). Upper beak depth and mandible width are significantly correlated in the Pom × Scan F_2_ population (R^2^ = 0.25, p = 1.65e-08): F_2_ individuals with deeper upper beaks tend to have wider mandibles (Supplemental Figure 6).

#### QTL controlling single linear dimensions

Finally, we identified three additional major-effect QTL associated with variation in linear measurements of the braincase and mandible. QTL on LG2 (LOD = 5.6, PVE = 19.8%), LG5 (LOD = 4.7, PVE = 16.9%), and LG10 (LOD = 5.0, PVE = 18.2%) are significantly associated with braincase length, braincase width, and mandible length, respectively (Supplemental Figures 5-6, Supplemental Tables 5-7). Taken together, our whole-genome scans revealed a set of seven major-effect QTL associated with linear measurements of the head skeleton that each explain 17-25% of the total phenotypic variance. We identified significant correlations between linear measurements of the same structure (e.g., upper beak width and depth) and of different structures (e.g., upper beak depth and mandible width); therefore, in some cases, regulation of multiple axes of craniofacial variation is coordinated by a single genomic locus.

### Geometric morphometric analyses of craniofacial shape variation

Linear measurements provide a simple description of some of the major axes of shape variation, but do not fully capture the complex 3D nature of the skull and mandible. We therefore used geometric morphometric methods (Zelditch *et al*. 2012; Adams *et al*. 2013) to analyze 3D shape variation by dividing the head into two substructures: (1) upper beak and braincase (UBB, 49 landmarks), and (2) lower beak or mandible (MAN, 24 landmarks). We assessed UBB and MAN shape integration by performing a two-block partial least squares (2B-PLS) analysis, which demonstrated that the main axis of integration (PLS1) is craniofacial curvature (r-PLS: 0.81, p < 0.001, Supplemental Figure 8A). In both substructures, allometry represents a small but significant component of shape variation: UBB and MAN shape are significantly positively associated with their respective centroid size (UBB R^2^ = 0.109, p < 0.001; MAN R^2^ = 0.069, p < 0.001); birds with larger head skeletons have a straighter, longer UBB and wider MAN (Supplemental Figure 8A-C). Allometry is an evolutionarily important associate of shape (De Beer 1940; Alberch *et al*. 1979; Hallgrímsson *et al*. 2019); however, we focused our further analyses on non-allometric shape variation within the Pom × Scan F_2_ population by using the residuals from the shape ∼ centroid size regression.

#### Upper beak and braincase (UBB) shape variation

Principal components analysis (PCA) demonstrated that the first 17 UBB PCs contribute to 90% of non-allometric shape variation in the Pom × Scan F_2_ population (Figure 3A). The first two UBB PCs account for ∼41% of total shape variation (Figure 3A). The principal axis of UBB shape variation (PC1, 30.11% of shape variation) represents variation in curvature along the entire length of the UBB anterior-posterior axis (Figure 3C, Supplemental Movie 1) and defines the most conspicuous difference between the craniofacial skeletons of the Pom and Scan founder breeds (Figure 1A-D). Within the PC1 morphospace, most F_2_ individuals are constrained by the cross founders, but cluster closer to the Pom founder than the Scan founder (Figure 3B).

**Figure 3.**
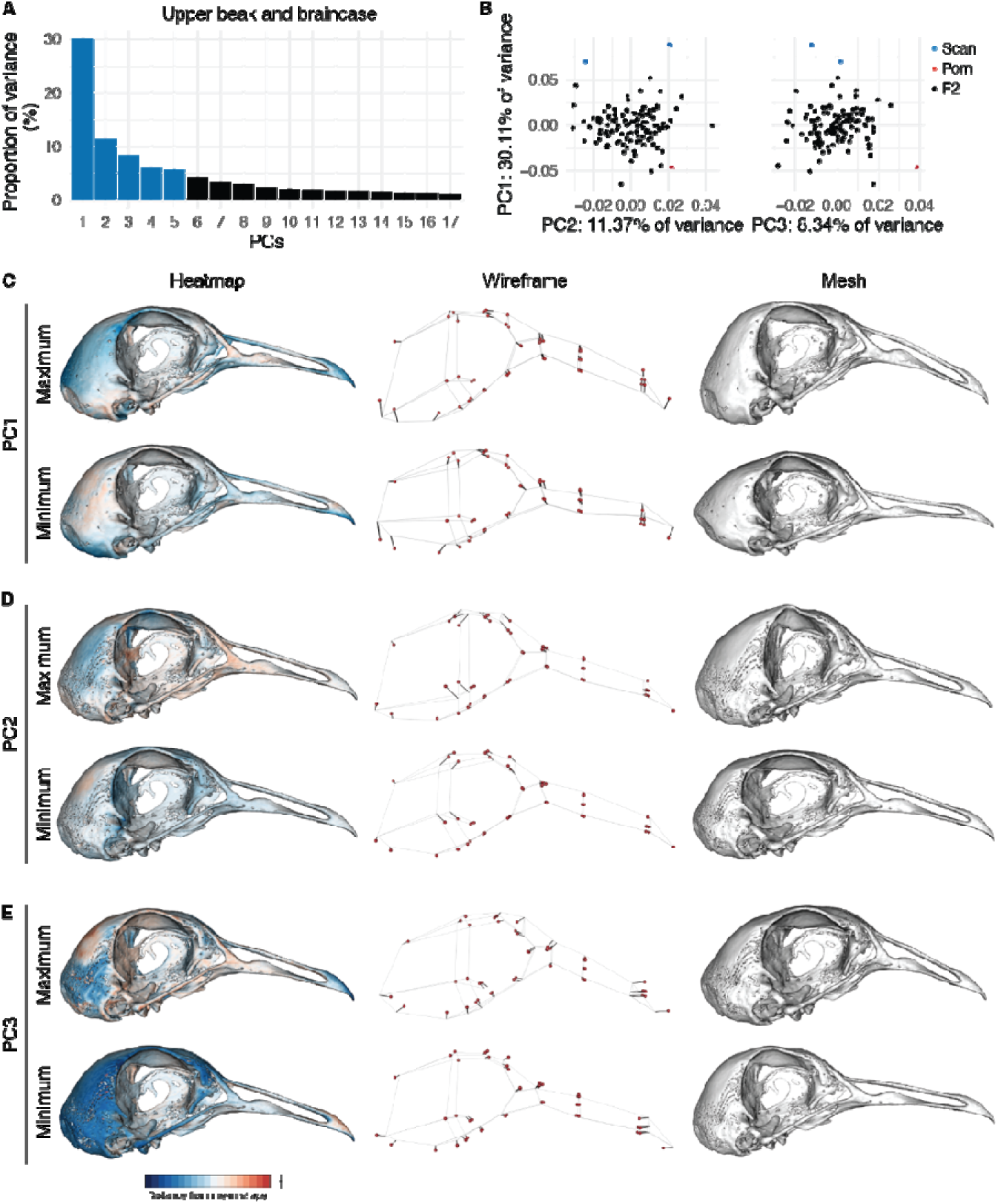
Upper beak and braincase (UBB) shape variation in the Pom × Scan F_2_ population. (A) Principal components (PCs) that collectively explain 90% of UBB shape variation. PCs that account for more than 5% of variation are indicated in blue. (B) PCA plots of PC1 vs. PC2 (left) and PC1 vs. PC3 (right). Founders are highlighted in blue (Scan) and red (Pom), F_2_ birds are denoted in black. (C-E) Visualizations of PC1 (C), PC2 (D), and PC3 (E) minimum and maximum shapes in three ways: heatmaps displaying distance from mean shape (left), wireframes showing displacement of landmarks from mean shape (center), and warped meshes (right). For wireframes and meshes, shape changes are magnified to aid visualization: 1.5x for PC1, 2x for PC2, 3x for PC3.

While PC1 incorporates landmarks from the entire UBB, PC2 (11.37% of UBB shape variation) is defined almost exclusively by variation in braincase shape (Figure 3D). The UBB PC2 axis describes the transition from a wide and shallow braincase (negative PC2 score) to a narrow and deep braincase (positive PC2 score; Figure 3D, Supplemental Movie 2). PC3-PC5 each account for 5-10% of UBB shape variation and describe complex 3D shape changes that involve landmarks from the upper beak and braincase (Figure 3E, Supplemental Figure 9, Supplemental Movies 3-5).

#### Mandible (MAN) shape variation

In the Pom × Scan F_2_ population, 90% of MAN shape is described by the first 13 PCs (Figure 4A). The first three PCs each describe >10% of variation and collectively account for ∼60% of total shape variation (Figure 4A). MAN PC1 (29.53% of total variation) describes a concomitant change in width and curvature, which results from displacement of both anterior and posterior landmarks (Figure 4C, Supplemental Movie 6). Unlike UBB PC1, MAN PC1 morphospace is not constrained by the cross founders: many F_2_ individuals have higher PC1 scores (narrower/straighter mandibles) than the founders (Figure 4B, Supplemental Figure 10).

**Figure 4.**
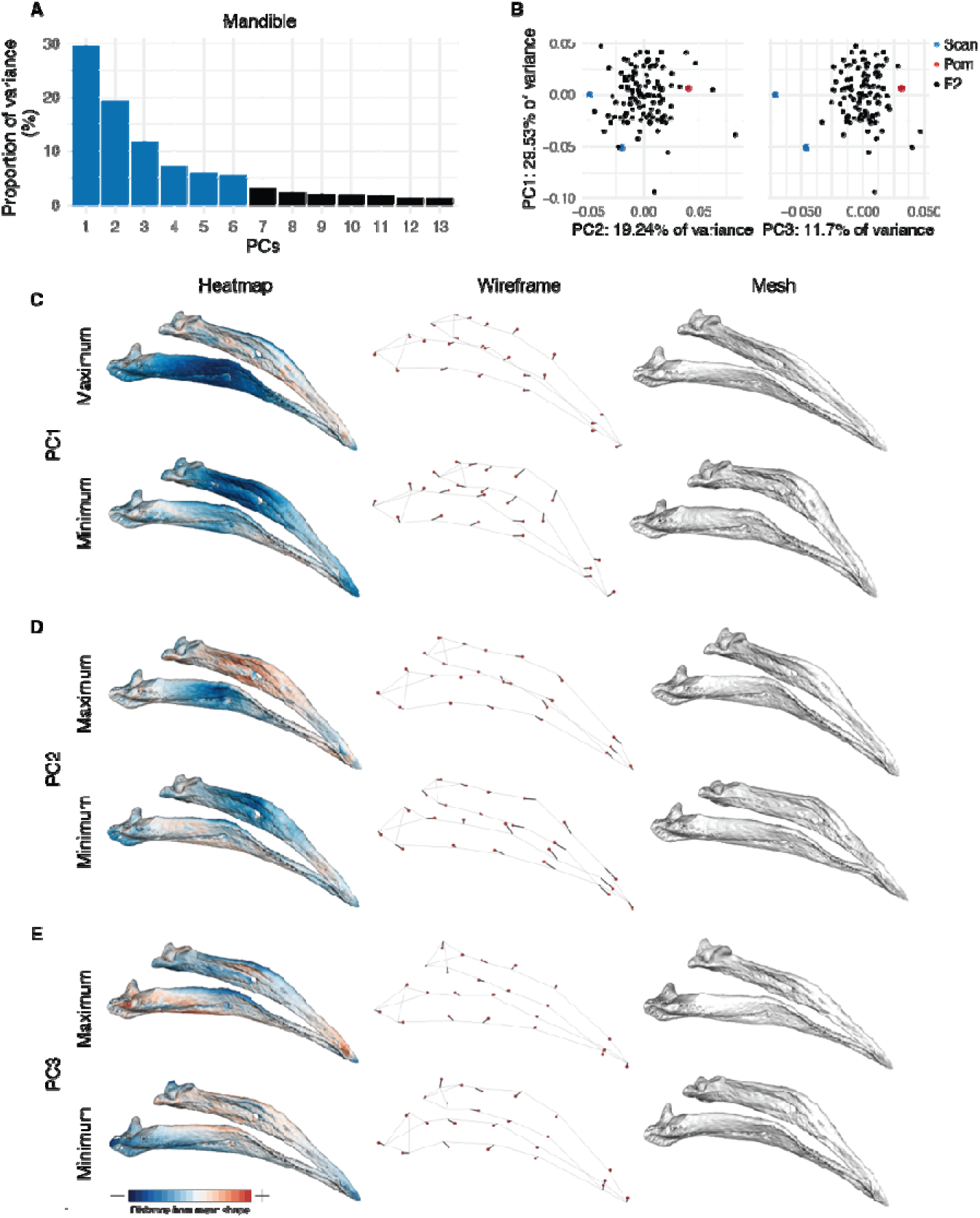
Mandible (MAN) shape variation in the Pom × Scan F_2_ population. (A) Principal components (PCs) that collectively explain 90% of MAN shape variation. PCs that account for more than 5% of variation are indicated in blue. (B) PCA plots of PC1 vs. PC2 (left) and PC1 vs. PC3 (right). Founders are highlighted in blue (Scan) and red (Pom), F_2_ birds are denoted in black. (C-E) Visualizations of PC1 (C), PC2 (D), and PC3 (E) minimum and maximum shapes in three ways: heatmaps displaying distance from mean shape (left), wireframes showing displacement of landmarks from mean shape (center), and warped meshes (right). For wireframes and meshes, shape changes are magnified to aid visualization: 1.5x for PC1 and PC2, 2x for PC3.

Positive scores for MAN PC2 (19.24% of variation) describe a narrowing at the center of the mandible and an elongation of the anterior mandible (Figure 4D, Supplemental Movie 7). PC3 (11.7% of variation) defines rotation in the posterior portion of the mandible that results in both increased posterior mandible width and reduced curvature along the entire length of the mandible in individuals with positive PC3 scores (Figure 4E, Supplemental Movie 8). PC4-6, which each account for 5-10% of total MAN variation, describe complex shape changes that affect aspects of mandible width (PC4, Supplemental Figure 11, Supplemental Movie 9), height (PC5, Supplemental Figure 12, Supplemental Movie 10), and curvature (PC6, Supplemental Figure 13, Supplemental Movie 11).

#### QTL associated with three-dimensional shape of the UBB

Next, we used the scores from the UBB and MAN PCs that explain >5% of total shape variation (PC1-5 for UBB, PC1-6 for MAN) to scan for QTL associated with shape variation. We identified four QTL associated with variation in UBB shape (summarized in Table 1). The UBB PC2 LOD support interval is a 17.3-Mb region that contains 171 genes, of which 146 are expressed during pigeon craniofacial development (Figure 5, Supplemental Table 8). F_2_ individuals homozygous for the Pom allele have higher UBB PC2 scores (taller, narrower braincases) than Scan homozygotes (Figure 5D), consistent with the shapes of the founders.

The UBB PC3 interval is a 1.3-Mb region that contains only 4 genes (*GAB3*, *SMARCA1*, *TENM1*, *SH2D1A*), all of which are expressed during pigeon craniofacial development (Supplemental Figure 14, Supplemental Table 9). In mouse embryos, *Gab3* and *Smarca1* are expressed in the first branchial arch (Brunskill *et al*. 2014), but their role in craniofacial development remains unknown. For UBB PC3, Pom homozygotes have lower scores (smaller braincase and longer, straighter upper beak) than Scan homozygotes, consistent with the result that the Pom founder sets the lower limit of the UBB PC3 morphospace (Figure 3B).

We identified two major-effect QTL associated with UBB PC4 on LG10 and LG11 (Supplemental Figure 15). The 10.2-Mb (LG10) and 16.0-Mb (LG11) intervals respectively contain 45 and 177 genes that are expressed during pigeon craniofacial development (Supplemental Figure 15, Supplemental Tables 10-11).

#### QTL associated with three-dimensional shape of the MAN

We also identified four QTL associated with MAN shape variation (summarized in Table 1). The LOD support intervals for the two MAN PC3 QTL encompass 1.9-Mb and 7.2-Mb genomic regions that contain 21 and 31 expressed genes, respectively (Figure 6B-C,E-F, Supplemental Tables 12-13). Notably, the LG2 interval includes the entire *HOXA* gene cluster. *HOXA2* is expressed during pigeon craniofacial development (Supplemental Table 12) and serves essential and evolutionarily-conserved roles in hindbrain, neural crest, and craniofacial patterning (Parker *et al*. 2018).

For MAN PC4, we identified a 1.4-Mb interval that contains 21 genes that are expressed during pigeon craniofacial development, including *FGF18* (Supplemental Figure 11, Supplemental Table 14). In mouse embryos, *Fgf18* functions in a molecular circuit with *Foxf* and *Shh* to regulate craniofacial development in mice (Xu *et al*. 2016; Yue *et al*. 2020).

Finally, the MAN PC5 LOD support interval is 0.54 Mb in length and includes 6 expressed genes (*ATG7*, *VGLL4*, *TAMM41*, *SYN2*, *TIMP4*, *PPARG*), none of which are known to contribute to craniofacial development (Supplemental Figure 12, Supplemental Table 15). In summary, we identified eight major-effect QTL that regulate 3D UBB and MAN shape variation, some of which contain genes with known roles in craniofacial development in other species, and others that do not.

#### Multi-locus QTL models describe major axes of Pom × Scan craniofacial shape variation

Our initial one-dimensional scans for major-effect QTL did not identify significant loci associated with UBB or MAN PC1. We predict this may be because, even after parsing skull and jaw shape variation into its component parts (PCs), UBB and MAN PC1 still describe highly complex 3D shape changes that likely have a polygenic basis. Although one-dimensional scans can detect multiple QTL (Broman *et al*. 2003), it is possible that PC1 shape is regulated by the combined action of many minor-effect QTL that we are underpowered to detect. Therefore, as an alternative strategy, we implemented multi-locus modeling and identified sets of 11 and 16 minor-effect QTL associated with UBB and MAN PC1 shape variation, respectively (Supplemental Tables 16 and 17). Although the multi-locus models suggest that each QTL set accounts for almost all of UBB and MAN PC1 shape variation (92.2% and 99.1%, respectively), additional undetected QTL might also contribute to UBB and MAN PC1 shape regulation, as estimated QTL effects are often biased upward, especially in relatively small mapping populations (Xu 2003).

## Discussion

Domestic species are remarkable repositories of phenotypic diversity (Darwin 1868; Andersson 2001; Rimbault and Ostrander 2012; Sánchez-Villagra *et al*. 2016). Unlike distantly related species with highly divergent phenotypes, breeds and strains of the same species – including those with radically different craniofacial traits – are interfertile, making genetic crosses and genomic comparisons experimentally tractable. Here, we used pigeon breeds with distinctive traits to map the genetic architecture of size and shape changes in the upper beak, braincase, and mandible. Overall, our results show that in pigeons, skull and jaw morphology has a complex genetic architecture, consistent with analyses of craniofacial shape in wild birds and other vertebrates (Albertson *et al*. 2003, 2005b; Schoenebeck *et al*. 2012; Pallares *et al*. 2015; Shaffer *et al*. 2016; Claes *et al*. 2018; Xiong *et al*. 2019; Yusuf *et al*. 2020; Katz *et al*. 2020).

### Coordinated and independent control of craniofacial traits

We identified 15 major-effect QTL associated with variation in skull and jaw shape in a pigeon F_2_ intercross (Figure 7). The QTL support intervals are dispersed across autosomes and the Z-chromosome, collectively span 117 Mb (∼10%) of the pigeon genome, and include 1104 genes. We measured skull and jaw shape using two methods – linear measurements and 3D shape – and found that QTL associated with variation in linear and 3D shape of the same structures did not overlap (Figure 7). Consistent with this finding, the 3D shape changes we quantified were not driven by changes in a single linear measurement, but were instead complex shape changes involving coordinated displacement of many landmarks. For the most part, skull and jaw shape QTL also did not overlap (Figure 7). Likewise, evidence from other species demonstrates that the vertebrate upper and lower jaws are largely modular structures that can evolve independently under separate genetic control. This genetic and developmental modularity, in turn, might facilitate the semi-independent evolutionary diversification of jaw and skull structures (Stockard and Johnson 1941; Drake and Klingenberg 2010; Parsons *et al*. 2011, 2018; Fish *et al*. 2011; Klingenberg 2014; Fish 2016; Felice and Goswami 2018; Bardua *et al*. 2019).

**Figure 5.**
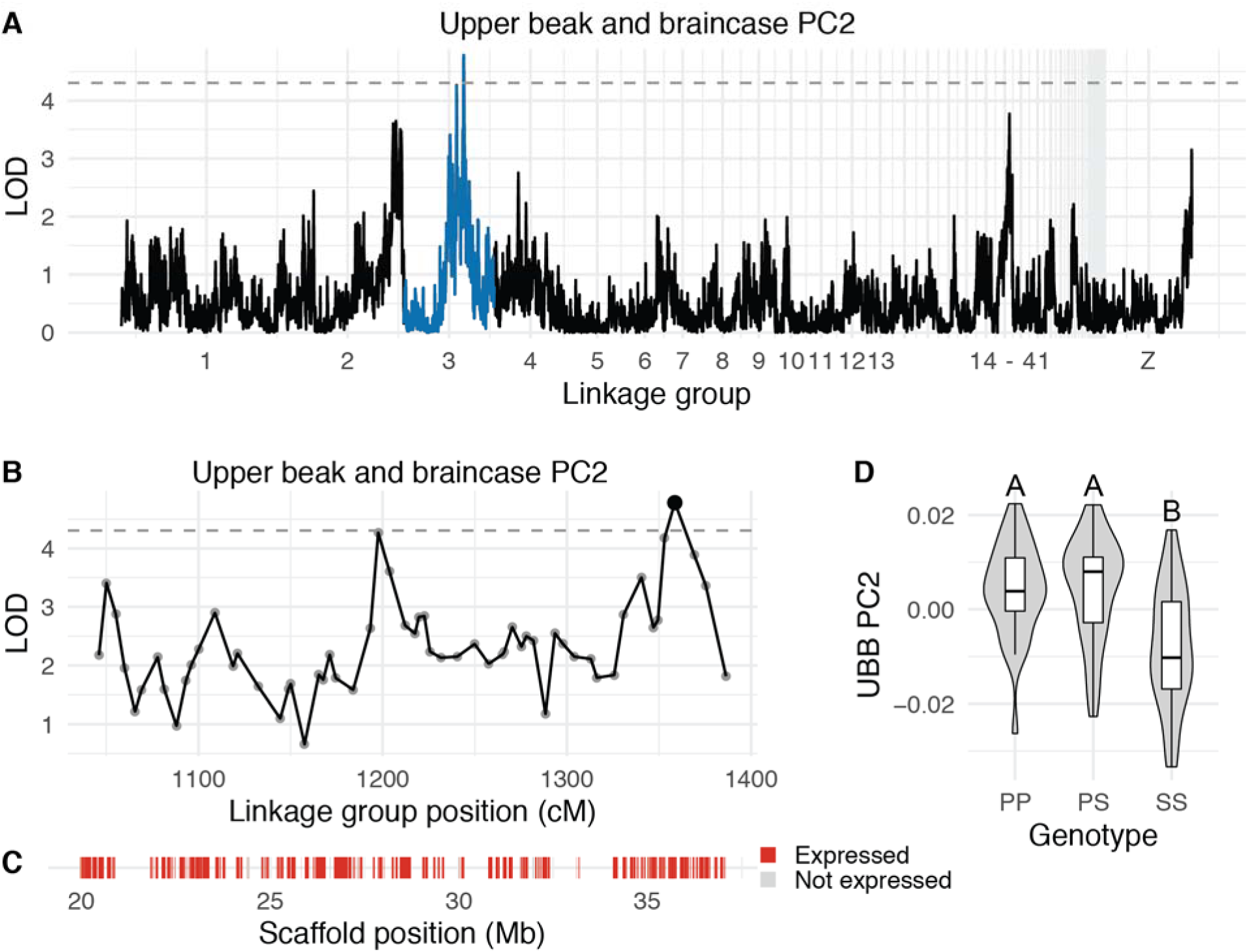
QTL associated with UBB PC2. (A) Genome-wide QTL scan for UBB PC2. Dashed horizontal line indicates 5% genome-wide significance threshold and linkage groups with significant QTL peaks are highlighted in blue. (B) LOD support interval for UBB PC2 QTL scan. Dots indicate linkage map markers; the larger black dot highlights the peak marker that was used to estimate QTL effects. (C) Genes located within LOD support interval, color coded based on expression status in HH29 facial primordia. (D) QTL effect plot for UBB PC2. Letters denote significance groups, p-values determined via Tukey test: PP vs. SS = 6.4e-04, PS vs. SS = 3.1e-05. P = allele from Pom founder, S = allele from Scan founder.

**Figure 6.**
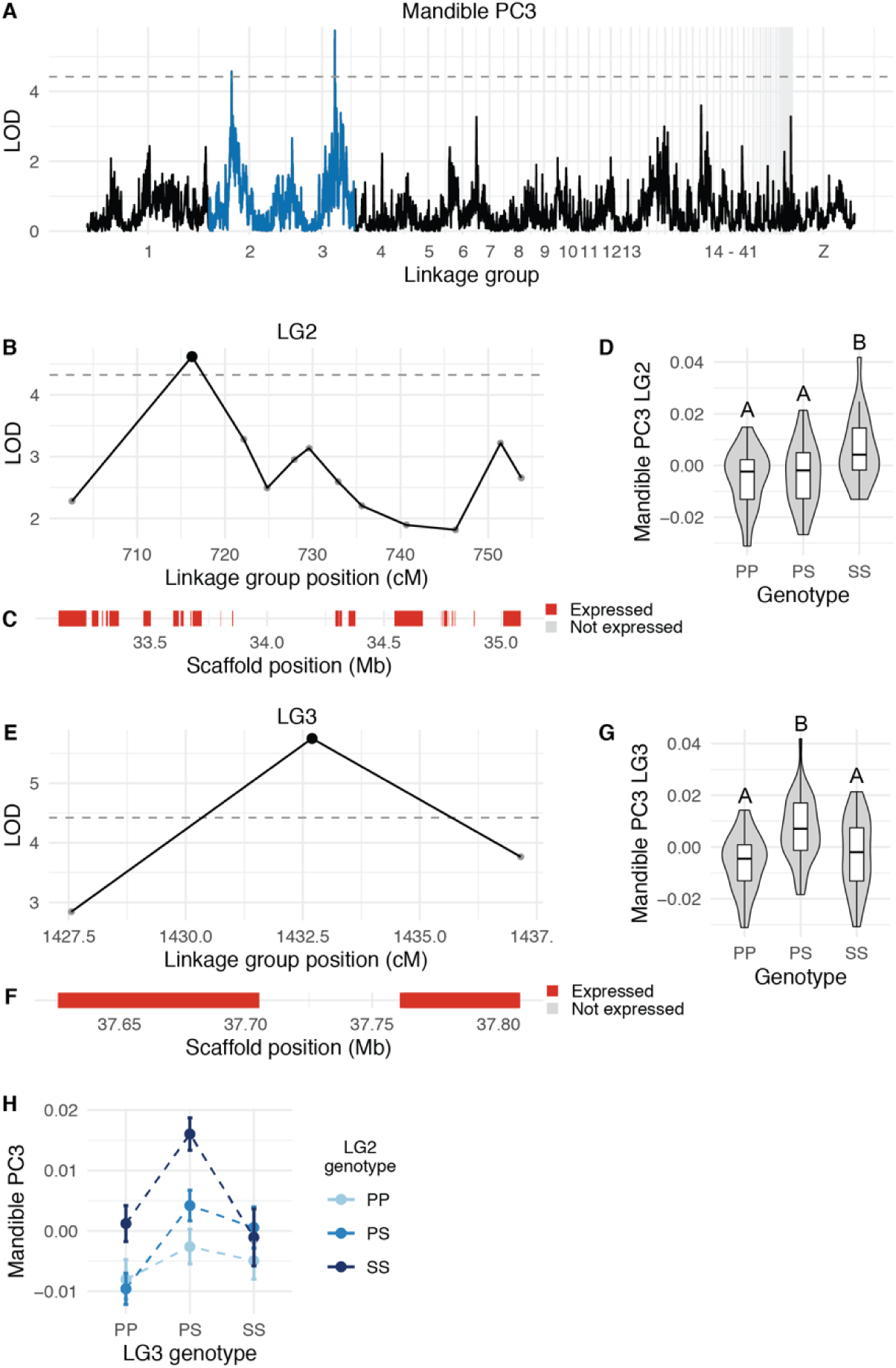
QTL associated with MAN PC3. (A) Genome-wide QTL scan for MAN PC3. Dashed horizontal line indicates 5% genome-wide significance threshold, and linkage groups with significant QTL peaks are highlighted in blue. (B) LOD support interval for MAN PC3 QTL on linkage group 2. Dots indicate linkage map markers; the larger black dot highlights the peak marker that was used to estimate QTL effects. (C) Genes located within LOD support interval, color coded based on expression status in HH29 facial primordia. (D) Effect plot for MAN PC3 QTL on LG2. Letters denote significance groups, p-values determined via Tukey test: PP vs. SS = 1.2e-04, PS vs. SS = 2.1e-03. (E) LOD support interval for MAN PC3 QTL on LG3. (F) Genes located within LG3 QTL. (G) Effect plot for QTL on LG3. Letters denote significance groups, p-values: PP vs. PS = 2.3e-05, PS vs. SS = 1.2e-02. (H) Interaction plot for MAN PC3 QTL on LG2 and LG3. P = allele from Pom founder, S = allele from Scan founder.

**Figure 7.**
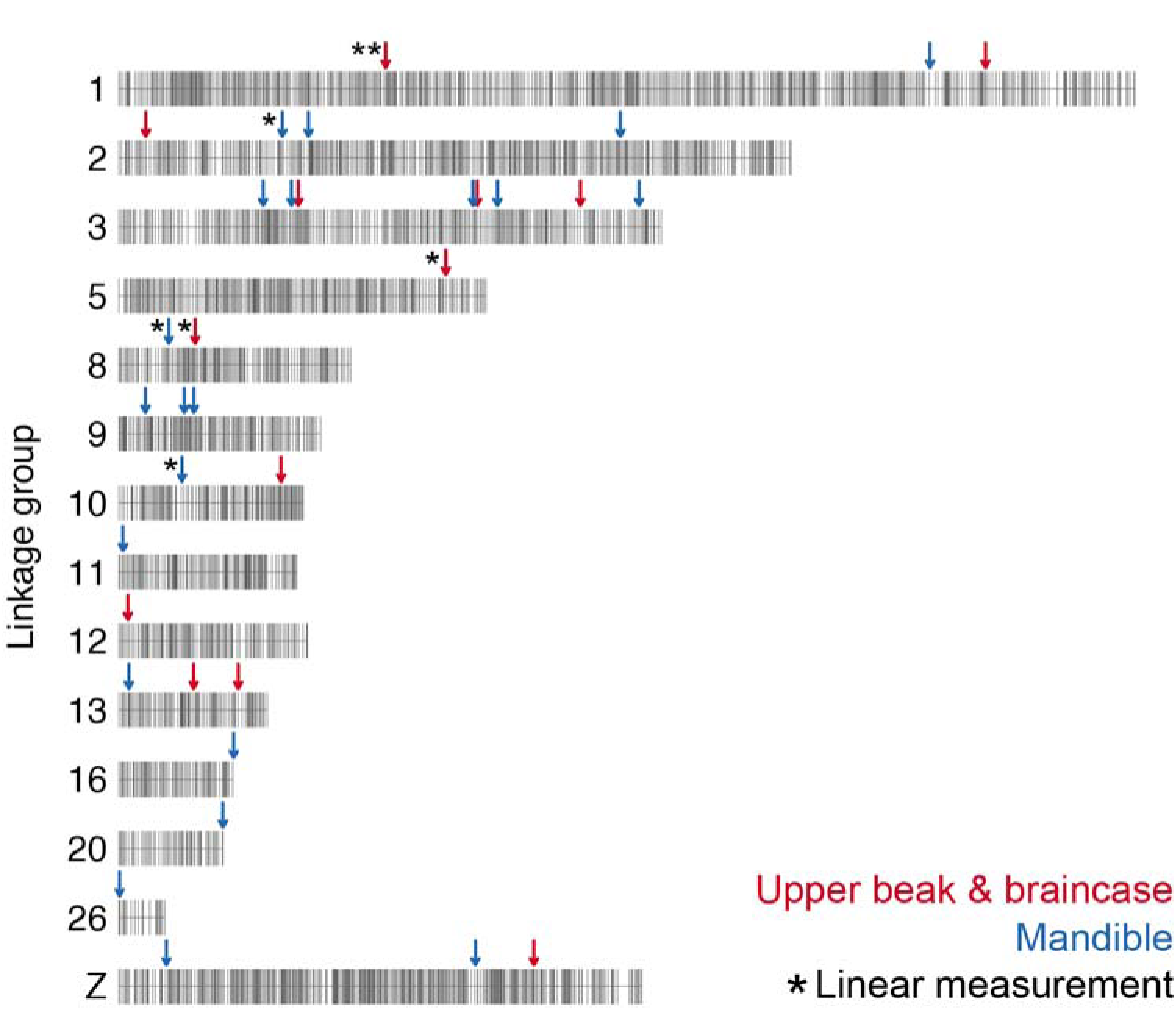
Summary of QTL associated with craniofacial shape in the Pom × Scan F_2_ population. Only the linkage groups harboring significant QTL are displayed. Markers are indicated by vertical gray lines. Approximate positions of QTL peaks are labeled with arrows; red and blue arrows mark QTL associated with UBB or MAN shape, respectively. Linear measurement QTL are indicated by asterisks to the left of the corresponding arrow; QTL without asterisks are associated with 3D shape changes.

Our QTL mapping experiments identified a set of genomic regions associated with craniofacial variation, but we currently do not know if these loci are specific to the Pomeranian Pouter and Scandaroon breeds, or if we have uncovered loci that broadly regulate craniofacial morphogenesis across pigeons, birds, or vertebrates. QTL mapping provides a powerful and direct link between genotype and phenotype but is also inherently limited because a mapping experiment can only assay genetic variation within a genetic cross, rather than survey genetic and morphological variation across the entirety of a species.

### Craniofacial curvature in pigeons

One of our principal goals was to identify genetic regulators of beak curvature. Our geometric morphometric analyses confirmed that craniofacial curvature was indeed the predominant axis of variation in the Pom × Scan F_2_ population. One unexpected finding from the geometric morphometric analyses is that within the UBB, beak curvature does not occur in isolation, but instead is linked to braincase curvature in a consistent and predictable manner (Figure 3C and Supplemental Movie 1). UBB and MAN curvature are also morphologically integrated (Supplemental Figure 8A), suggesting that coordinated genetic programs contribute to development of the upper and lower beak. However, we did not identify QTL that regulate both UBB and MAN shape. It is possible that shared QTL are either beyond our limit of detection in the Pom × Scan cross, or that distinct UBB and MAN QTL harbor genes that belong to a common genetic program.

Along the UBB PC1 (curvature) axis, we found that many Pom × Scan F_2_ progeny approach or exceed the shape of the Pom founder, but never the Scan founders. This finding suggests that the straight-beaked Pom phenotype (closer to the ancestral condition) results from a variety of genotype combinations at different loci, but the extreme craniofacial curvature that defines the Scan breed probably requires the combined action of specific alleles at many loci. The Scandaroon is one of the oldest breeds of domestic pigeon (Levi 1986); millennia of artificial selection likely fixed a polygenic program to consistently produce the breed-defining enlarged and curved beak. Our F_2_ population was probably not big enough to have an appreciable (or any) number of offspring with the right allelic combinations to recapitulate the Scan craniofacial phenotype.

### Complex genetic architecture of an exaggerated craniofacial trait

The enlarged, curved craniofacial skeleton of the Scandaroon breed is a spectacular example of an exaggerated trait (an elaboration of an ancestral trait). To date, our understanding of the genetic basis of exaggerated traits remains relatively limited relative to trait reduction or loss. The pigeon craniofacial skeleton offers a unique opportunity to compare trait exaggeration and reduction: in addition to the exaggerated beak morphology of the Scandaroon breed, many breeds have dramatically reduced beaks (e.g., breeds from the Owl and Tumbler families). In our recent investigation of the genetic basis of the short beak phenotype in pigeons, we found that a single major-effect locus explains the majority of variation in beak reduction (Boer *et al*. 2021).

Here, we tested the outcome of shuffling the genomes of two divergent pigeon breeds and found that, even in this relatively simple context, many genetic regions are involved in determining craniofacial exaggeration. The results of the Pom × Scan F_2_ intercross are consistent with findings from classical genetic experiments performed in pigeons over the last century (Christie and Wriedt 1924; Sell 2012), in which elaboration of beak size has a separate and more complicated genetic architecture than beak reduction. Our results are also consistent with studies of craniofacial genetics from diverse vertebrates; the prevailing model is that the genetic architecture of craniofacial variation is highly polygenic (Richmond *et al*. 2018; Yusuf *et al*. 2020). In humans, a multitude of genes encoding members of diverse molecular classes (e.g., cell adhesion and motility, signal transduction, transcriptional regulation, ribosome biogenesis) are implicated in both normal and pathogenic craniofacial variation (Shaffer *et al*. 2016; Claes *et al*. 2018; Weinberg *et al*. 2018; Richmond *et al*. 2018; Xiong *et al*. 2019).

Recent examples of trait exaggeration in other tissues, such as ornamental feathering in pigeons (Shapiro *et al*. 2013; Domyan *et al*. 2016) or fleshy snouts in cichlids (Concannon and Albertson 2015; Conith *et al*. 2018) show that morphological exaggeration can have a relatively simple genetic basis, in which a majority of the variation is explained by one or two genetic factors. In contrast, our results from the pigeon craniofacial skeleton suggest that multiple loci exert a substantial influence on beak elaboration.

## Materials and methods

### Animal husbandry and 3D imaging

All animal experiments, husbandry, and housing protocols for this study were approved by the University of Utah Institutional Animal Care and Use Committee (protocols 10-05007, 13-04012, and 19-02011).

An intercross between a male Pomeranian Pouter and two female Scandaroons was performed to generate 131 F_2_ offspring (Domyan *et al*. 2014, 2016). Cross founders and F_2_ individuals that survived to at least 6 months of age (n = 116) were euthanized and submitted to the University of Utah Preclinical Imaging Core Facility for micro-CT imaging. For each bird, a whole-body scan was performed on a Siemens Inveon micro-CT using the following parameters: voxel size = 94 μ, photon voltage = 80 kV, source current = 500 μA, exposure time = 200 ms. Scans were reconstructed using a Feldkamp algorithm with Sheep-Logan filter and a calibrated beam hardening correction. Of the F_2_ individuals that did not survive to maturity, 15 were used to construct the genetic map (see section on Genotyping and linkage map assembly).

### Surface model generation and landmarking

From the micro-CT image data, a substack that included the cranium was extracted from the whole-body DICOM file stack and saved in the NifTI format (*.nii) using ImageJ 1.52q. NifTI files were imported into Amira 6.5.0 software (Thermo Fisher Scientific) to generate a 3D surface model of the cranial skeleton. Using the threshold feature in Amira’s Segmentation Editor, the cranial skeleton was segmented from soft tissue. The resulting surface model was simplified and saved in the HxSurface binary (*.surf) format. Surface meshes were converted to the Polygon (Stanford) ASCII file format (*.ply) using i3D Converter v3.80 and imported into IDAV Landmark Editor v3.0 (UC Davis) for landmarking. An atlas of midline and bilateral Type 1 (defined by anatomy) and Type 3 (defined mathematically) landmarks on the braincase (29 landmarks), upper beak (20 landmarks), and mandible (24 landmarks) was developed using the pigeon atlas described in (Young *et al*. 2017) as a foundation. After landmarks were applied to 116 F_2_ individuals and the cross founders, the coordinates were exported as a NTsys landmark point dataset (*.dta) for geometric morphometric analysis.

### Morphometric analyses and shape change visualization

For each F_2_ individual and the cross founders, linear distances between sets of two landmarks (Supplemental Table 1) were measured in Landmark Editor. For each linear measurement, normal distribution within the F_2_ population was assessed using Shapiro-Wilk’s test in R v3.6.3 (R Core Team 2020). To account for differences in body size, each linear measurement was fit to a linear regression model (linear measurement ∼ body mass) and residuals were calculated in R. To compare residuals between sexes, a two-sided Wilcoxon test was implemented in R.

Geometric morphometric analyses were performed using the R package geomorph v3.3.1 (Collyer and Adams 2018, 2020; Adams *et al*. 2020). Briefly, the NTsys landmark point dataset was read in using the *readland.nts* function. The location of missing landmarks was estimated using the function *estimate.missing(method = “TPS”)*. We performed bilateral symmetry analysis via the function *bilat.symmetry(iter = 1)* and the symmetrical component of shape variation was extracted. After subsetting the data into two modules representing either upper beak and braincase (UBB) or mandible (MAN), we performed a Generalized Procrustes Analysis using the *gpagen* function. To analyze allometry, a linear model (shape ∼ centroid size) was fit using the *procD.lm* function and we used the residuals for analysis of allometry-free shape. We performed principal components analysis using the *gm.prcomp* function and analyzed integration using the *two.b.pls* function.

We visualized shape changes in geomorph and in the R package Morpho v2.8 (https://github.com/zarquon42b/Morpho). The geomorph function *plotRefToTarget* was used to generate wireframes. We generated surface mesh deformations, heatmaps, and movies in Morpho with the *tps3d*, *shade3d*, *meshDist*, and *warpmovie3d* functions. For all mesh-based visualizations, deformations were applied to a reference mesh. The reference mesh was created by warping a Pom × Scan F_2_ mesh to the mean shape.

### Genotyping and linkage map assembly

For cross founders and a subset of F_2_ individuals, we performed genotyping-by-sequencing (GBS) as previously described (Domyan *et al*. 2016). GBS libraries for an additional 20 F_2_ individuals, as well as supplemental libraries to improve coverage for 17 previously-sequenced individuals, were prepared and sequenced by the University of Minnesota Genomics Center. GBS libraries were sequenced on a NovaSeq 1×100 SP FlowCell. Target sequencing volume was ∼4.75 million reads/sample.

GBS reads were trimmed using CutAdapt (Martin 2011), then mapped to the Cliv_2.1 reference genome (Holt *et al*. 2018) using Bowtie2 (Langmead and Salzberg 2012). Genotypes were called using Stacks2 by running *refmap.pl* with the Pom and one of the two Scan founders designated as parents (Catchen *et al*. 2011, 2013). To account for the three-founder cross structure, we subsequently removed all markers where the genotypes of the two Scan founders differed; therefore, all alleles could be identified as originating from either the Pom or Scan founder breeds.

Genetic map construction was performed using R/qtl (www.rqtl.org; (Broman *et al*. 2003)). For autosomal markers, we eliminated markers showing significant segregation distortion (*p* < 0.01 divided by the total number of markers genotyped, to correct for multiple testing). We assembled and ordered sex-linked scaffolds separately, due to differences in segregation pattern for the Z chromosome. We identified Z-linked scaffolds by assessing sequence similarity and gene content between pigeon scaffolds and the Z chromosome of the annotated chicken genome assembly (Ensembl Gallus_gallus-5.0).

Pairwise recombination frequencies were calculated for all autosomal and Z-linked markers. We identified markers with identical genotyping information by using the *findDupMarkers* function, and then removed all but one marker in each set of duplicates. Within individual Cliv_2.1 scaffolds, markers were filtered by genotyping rate; to retain the maximum number of scaffolds in the final map, we performed an initial round of filtering to remove markers where fewer than 50% of birds were genotyped. Large scaffolds (> 40 markers) were subsequently filtered a second time to remove markers where fewer than 66% of birds were genotyped.

We used the R/qtl functions *droponemarker* and *calc.errorlod* to assess genotyping errors within individual scaffolds. Markers were removed if dropping the marker led to an increased LOD score, or if removing a non-terminal marker led to a decrease in preliminary linkage group length of >10 cM that was not supported by physical distance. Individual genotypes were removed if they showed an error LOD score >5 (Lincoln and Lander 1992). After these iterative rounds of filtering and quality control, we assembled final linkage groups from 3759 autosomal markers and 422 Z-linked markers using the parameters (max.rf 0.15, min.lod 6). Scaffolds in the same linkage group were manually ordered based on calculated recombination fractions and LOD scores.

### QTL mapping and LOD interval identification

We performed QTL mapping using R/qtl v1.46-2 (Broman *et al*. 2003). For each linear measurement residual and shape PC phenotype, we ran a single-QTL genome scan using the *scanone* function and Haley-Knott regression with sex as a covariate. For each phenotype, the 5% genome-wide significance threshold was calculated by running *scanone* with 1000 permutation replicates. A “major-effect QTL” was defined as any significant peak that was identified in a single-QTL genome scan. For phenotypes with significant QTL peaks, we calculated 1.5-LOD support intervals using the *lodint* function and estimated QTL effects via the *plotPXG* function. We compared phenotypic means in Pom × Scan F_2_ genotypic groups at peak markers via one-way ANOVA and Tukey Test for pairwise comparisons in R. For single-locus QTL, we calculated percent variance explained (PVE) using the *fitqtl* function.

To build multi-locus QTL models, two-dimensional genome scans were performed using the *scantwo* function. We identified candidate additive and interactive QTL using LOD thresholds lod.full = 9.1, lod.fv1 = 7.1, lod.int = 6.3, lod.add = 6.3, and lod.av1 = 3.3, as suggested by the R/qtl authors (Broman and Sen 2009). Multi-locus models were built using the *makeqtl*, *fitqtl*, and *refineqtl* functions. We identified genes within QTL intervals using a custom R script and visualized their locations using the R packages ggplot2 v3.3.0 (Wickham 2016) and gggenes v0.4.0 (https://github.com/wilkox/gggenes).

### RNA isolation, sequencing, and transcript quantification

Fertilized pigeon eggs were collected from Racing Homer (RH) and Oriental Frill (OF) breeding pairs and incubated to the equivalent of Hamburger-Hamilton stage 29 (HH29, embryonic day 6). We dissected the facial primordia (n = 5 from each breed) and stored the tissue in RNAlater (Thermo Fisher Scientific) at −80°C. We later extracted total RNA from each tissue sample using the RNeasy Mini Kit with RNase-Free DNAse Set and a TissueLyser LT instrument (Qiagen). RNA-sequencing libraries were prepared and sequenced by the High-Throughput Genomics and Bioinformatic Analysis Shared Resource at the University of Utah. RNA sample quality was assessed using the RNA ScreenTape Assay (Agilent) and sequencing libraries were prepared using the TruSeq Stranded mRNA Sample Prep Kit with oligo(dT) selection (Illumina). 125-cycle paired-end sequencing was performed on an Illumina HiSeq 2500 instrument (12 libraries/lane).

We assessed sequencing read quality with FastQC (Babraham Bioinformatics) and trimmed Illumina adapters with Cutadapt (Martin 2011). Reads were then aligned to the pigeon Cliv_2.1 reference assembly (Holt *et al*. 2018) and quantified using Salmon (Patro *et al*. 2017). Based on mean TPM (which was calculated from all samples), we characterized gene expression level as no expression/below cutoff (<0.5 TPM), low (0.5-10 TPM), medium (11-1000 TPM), or high (>1000 TPM), as described in the EMBL-EBI Expression Atlas (https://www.ebi.ac.uk/gxa/home).

## Acknowledgements

We are grateful to Nathan Young for generously sharing his time and expertise related to geometric morphometrics. We also thank Rich Schneider for thoughtful discussions and input on the project. We thank the organizer and instructors of the PR Statistics course on geometric morphometrics: Oliver Hooker, Dean Adams, Michael Collyer, and Antigoni Kaliontzopoulou. We are grateful to all past and present members of the Shapiro Lab, particularly Rebecca Bruders, Alexa Davis, Eric Domyan, Hannah Van Hollebeke, and Anna Vickrey for technical assistance and advice. We thank members of the Utah Pigeon Club and National Pigeon Association for sample contributions. We acknowledge the University of Utah Preclinical Imaging Core Facility, especially Tyler Thompson, for micro-CT imaging; the Center for High Performance Computing at the University of Utah for computing resources; the University of Utah High-Throughput Genomics Shared Resource for RNA library preparation and sequencing; and the University of Minnesota Genomics Core for GBS library preparation and sequencing.

## Data availability

RNA-sequencing datasets generated for this study have been deposited to the NCBI SRA database under BioProject PRJNA680754.

---

**Supplemental Figure 1.**
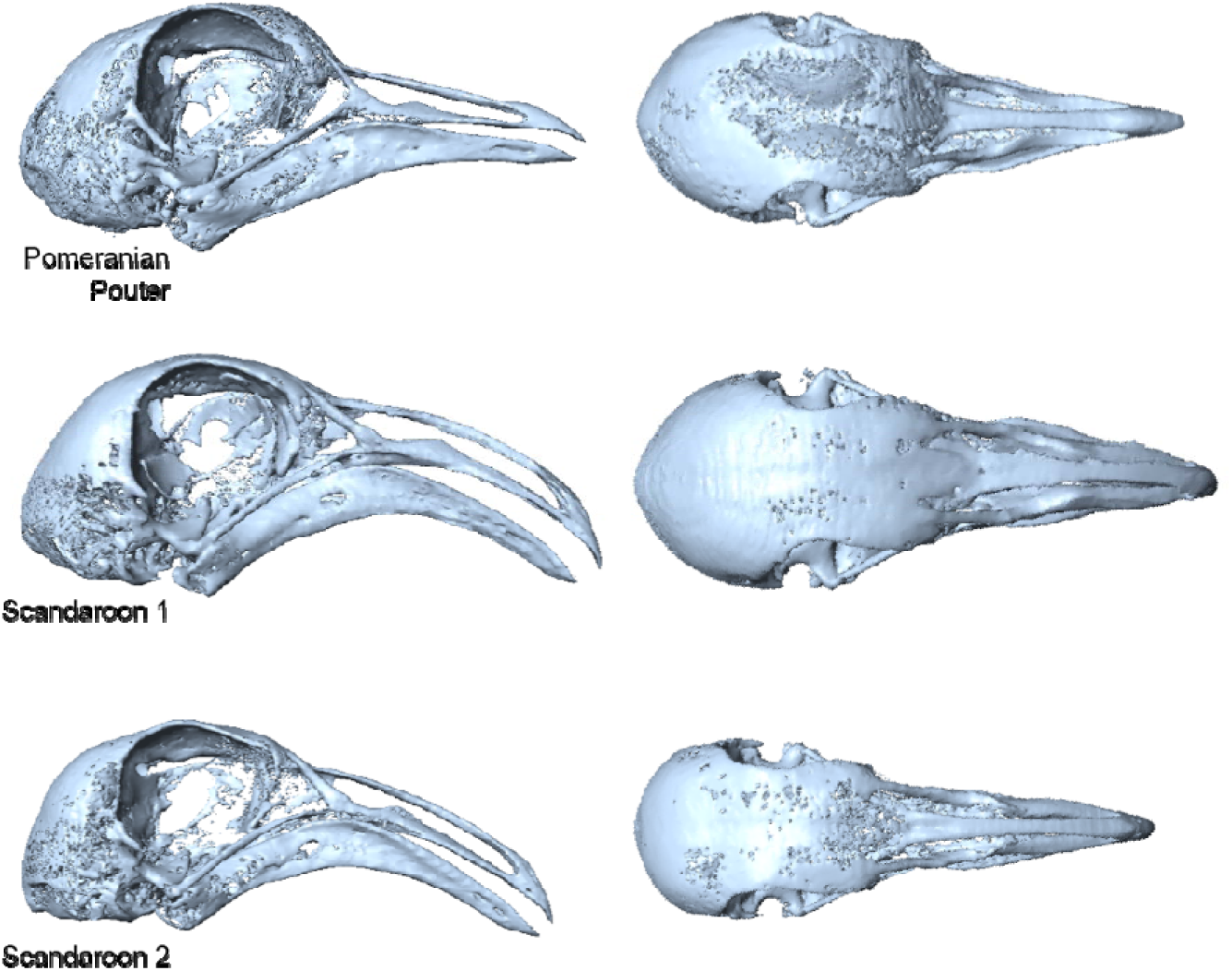
Surface models of the Pom × Scan founders. Lateral (left) and dorsal (right) views of the craniofacial skeleton of the male Pom and female Scan founders used to generate the F_2_ intercross.

**Supplemental Figure 2.**
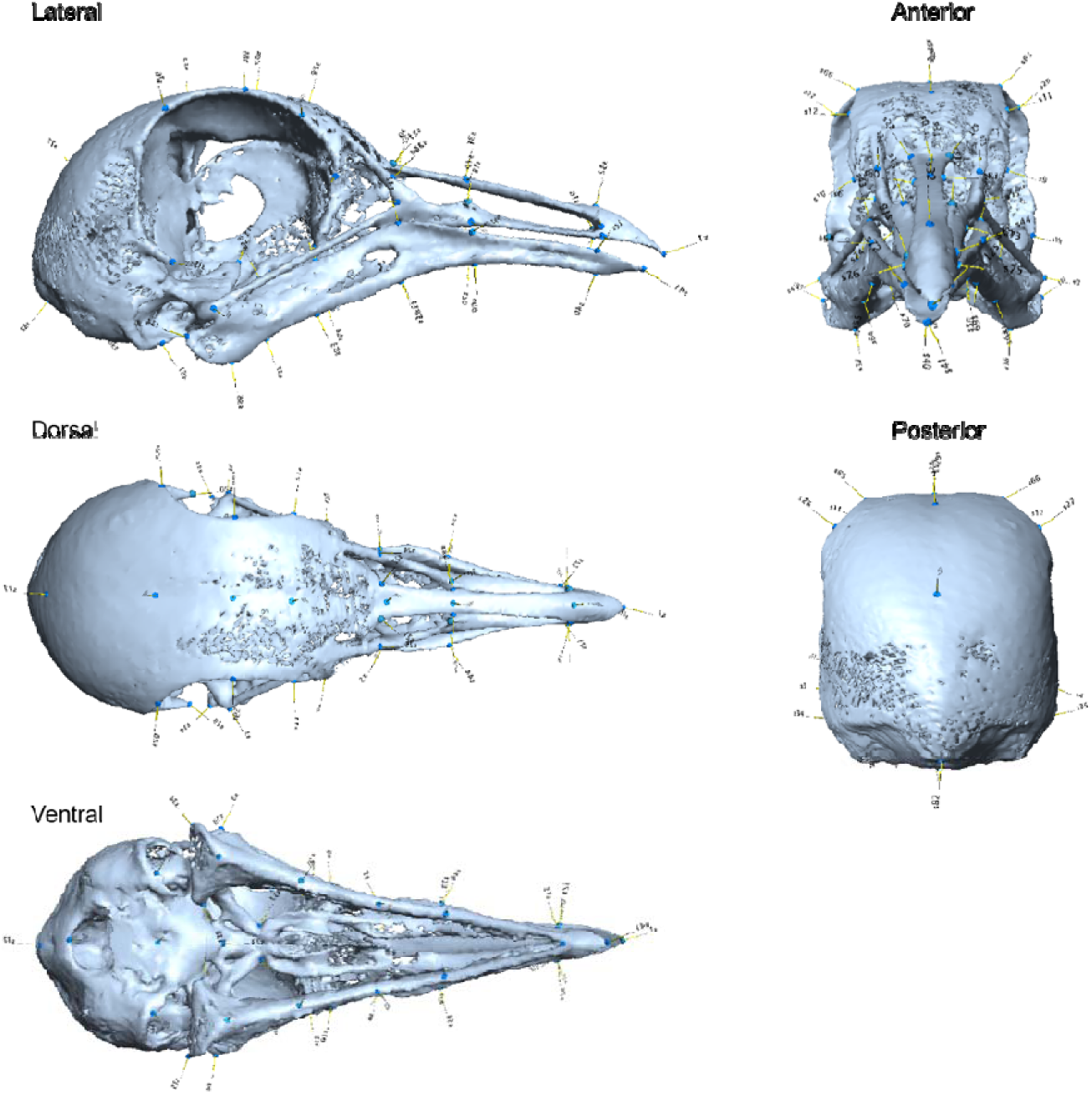
Pigeon craniofacial landmark atlas. Landmark positions are indicated by blue discs.

**Supplemental Figure 3.**
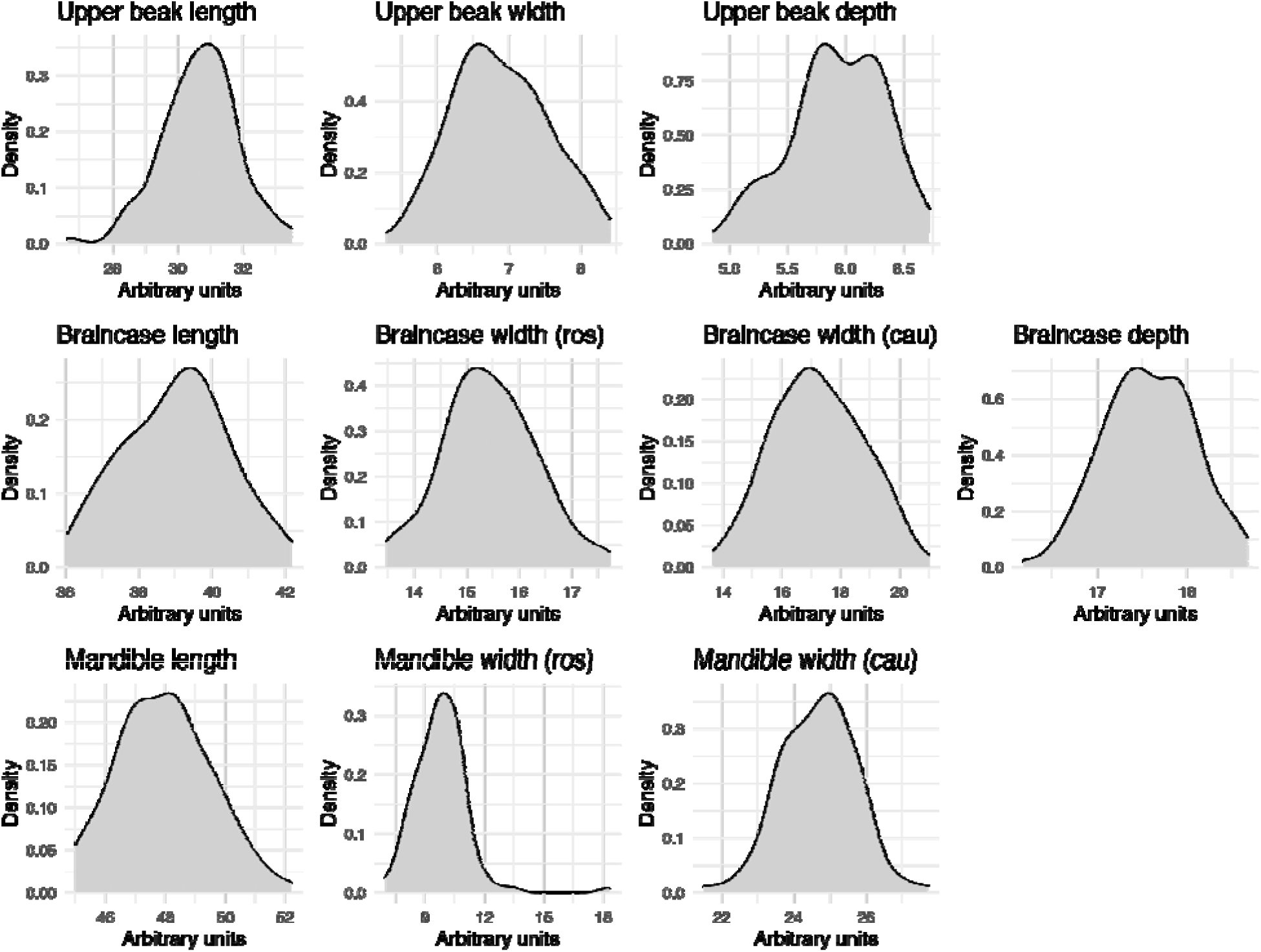
Distribution of 11 linear measurements in the Pom × Scan F_2_ population. With the exception of rostral mandible width, all linear measurements are normally distributed in the population (Shapiro-Wilk’s test, p > 0.05). For rostral mandible width, a single F_2_ individual is an outlier (MDS079, see Supplemental Figure 10) and causes a deviation from normality (Shapiro-Wilk’s p = 5.2e-09).

**Supplemental Figure 4.**
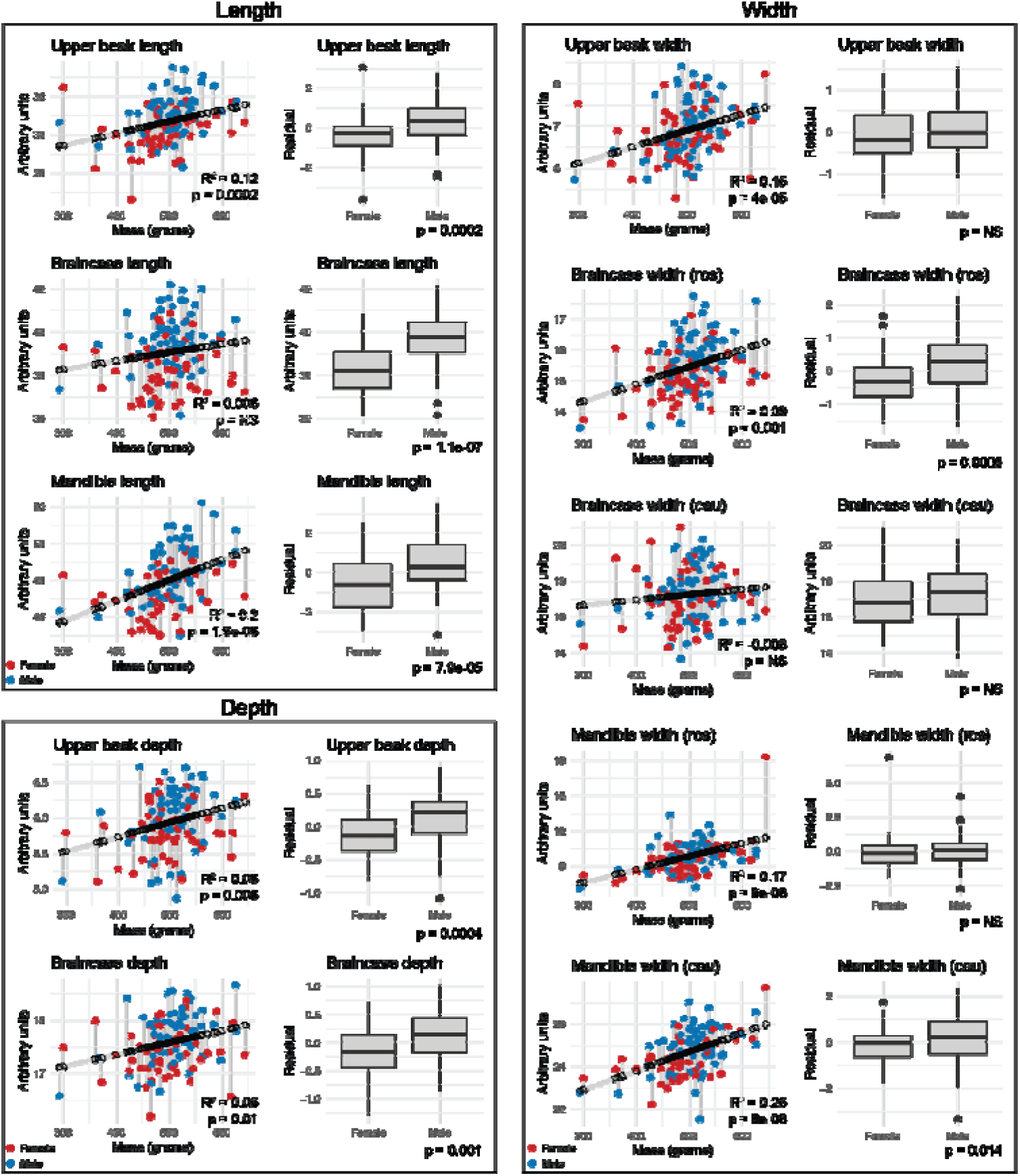
Linear regression of 11 craniofacial measurements on body mass. For all panels, linear measurement ∼ mass regression is displayed on the left with associated R^2^ and p-value are indicated in bottom right corner of each plot. Each dot represents raw measurement of an F_2_ individual, color-coded by sex (male = blue, female = red). Each raw measurement is connected to an open circle that indicates its predicted value; grey connecting lines correspond to residual value used for QTL mapping. In each panel, the boxplot on the right displays residual values by sex; outliers are indicated by black dots. Associated p-values are indicated in bottom right corner of each plot (two-sided Wilcoxon test).

**Supplemental Figure 5.**
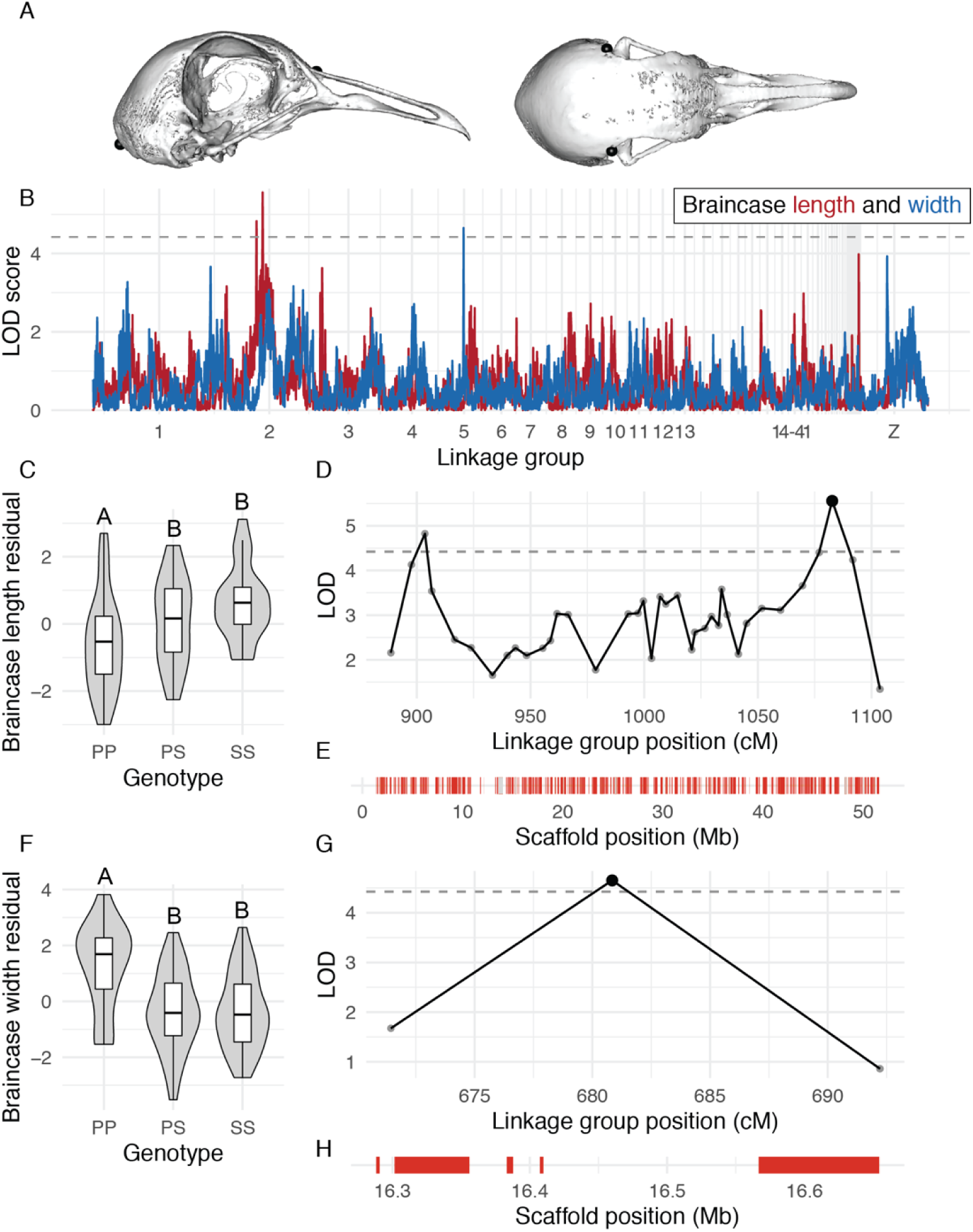
QTL associated with braincase length and width. (A) Landmark pairs used to measure braincase length (left) and width (right). (B) QTL scans for braincase length (red) and width (blue). Dashed horizontal lines denotes 5% genome-wide significance threshold. (C) Effect plot for braincase length QTL on LG2. Letters denote significance groups, p-values determined via Tukey test: PP vs. PS = 8.7e-03, PP vs. SS = 2.2e-04. (D) LOD support interval for braincase length on LG2. Dots indicate linkage map markers; the black dot highlights the peak marker that was used to estimate QTL effects. (E) Genes located within braincase length QTL LOD support interval, color coded based on if gene is expressed in HH29 facial primordia (red) or not expressed (gray). (F) Effect plot for braincase width QTL on LG5. Letters denote significance groups, p-values: PP vs. PS = 5.8e-05, PP vs. SS = 7.5e-05. (G) LOD support interval for braincase width QTL on LG5. (H) Genes located within braincase width QTL on LG5. P = allele from Pom founder, S = allele from Scan founder.

**Supplemental Figure 6.**
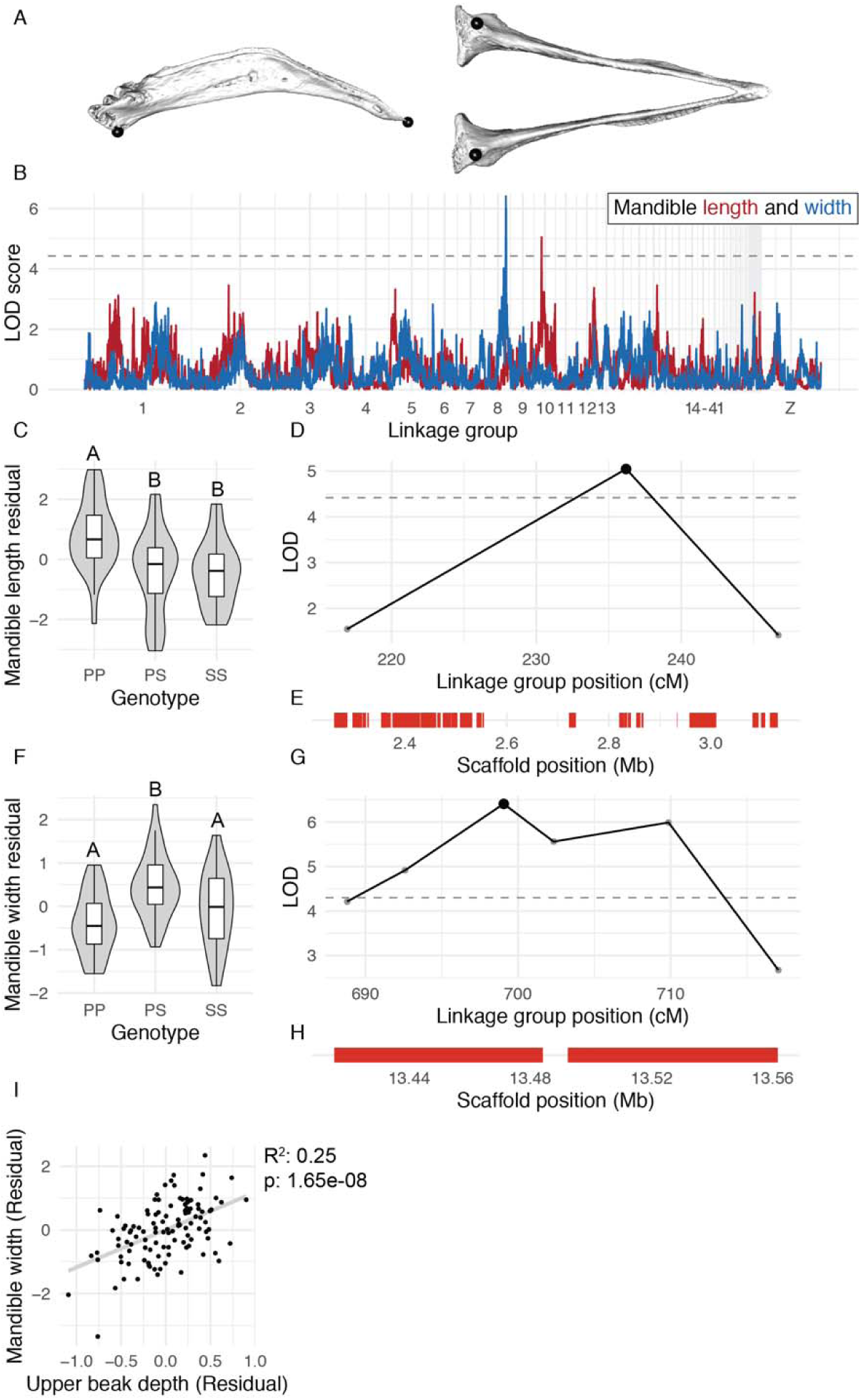
QTL associated with mandible length and width. (A) Landmark pairs used to measure mandible length (left) and width (right). (B) QTL scans for mandible length (red) and width (blue). Dashed horizontal lines denotes 5% genome-wide significance threshold. (C) Effect plot for mandible length QTL on LG8. Letters denote significance groups, p-values determined via Tukey test: PP vs. PS = 5.9e-04, PP vs. SS = 1.1e-03. (D) LOD support interval for mandible length QTL. (E) Genes located within mandible length QTL LOD support interval. (F) Effect plot for mandible width QTL on LG10. Letters denote significance groups, p-values determined via Tukey test: PP vs. PS = 2.7e-05, PS vs. SS = 1.7e-02. (G) LOD support interval for mandible width QTL. (H) Genes located within mandible width QTL. (I) Scatterplot of upper beak depth and mandible width residuals for all F_2_ individuals. P = allele from Pom founder, S = allele from Scan founder.

**Supplemental Figure 7.**
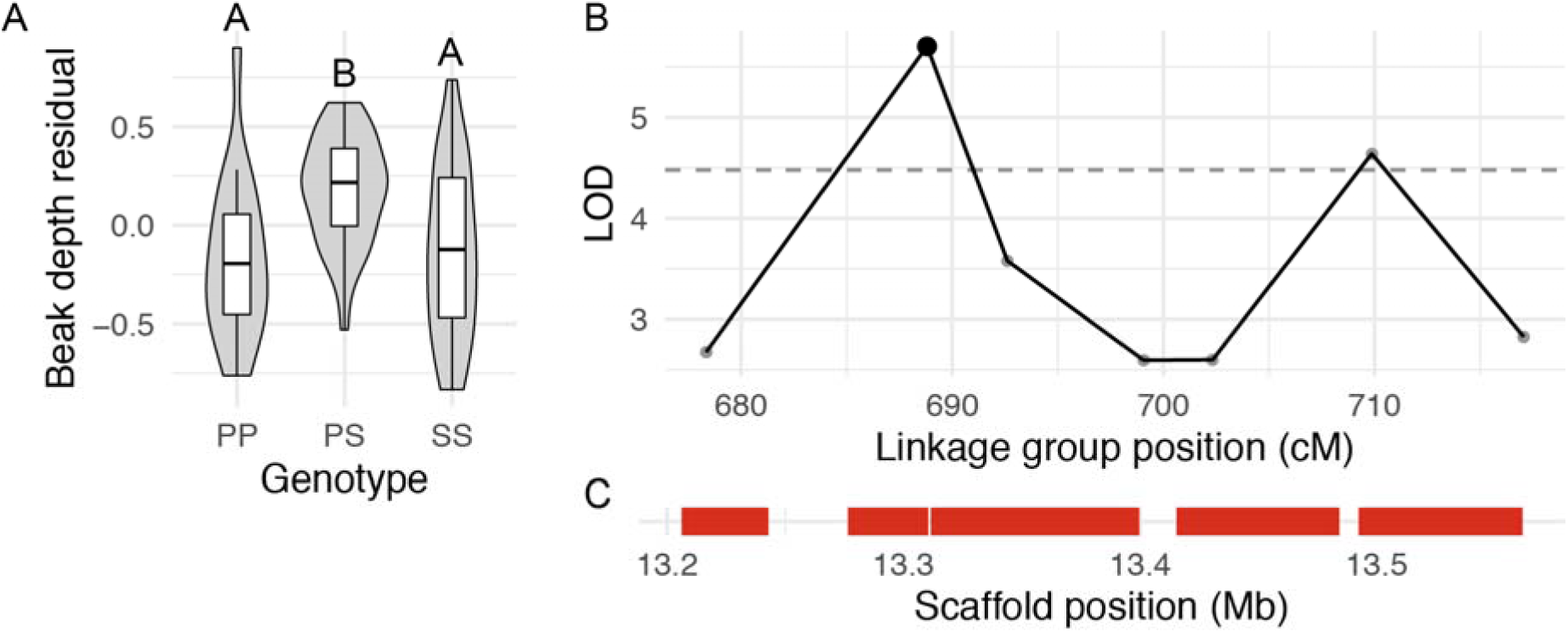
QTL on LG8 associated with upper beak width. (A) Effect plot for upper beak depth QTL. Letters denote significance groups, p-values determined via Tukey test: PP vs. PS = 3.9e-04, PS vs. SS = 1.7e-02. (B) LOD support interval. (C) Genes in interval. P = allele from Pom founder, S = allele from Scan founder.

**Supplemental Figure 8.**
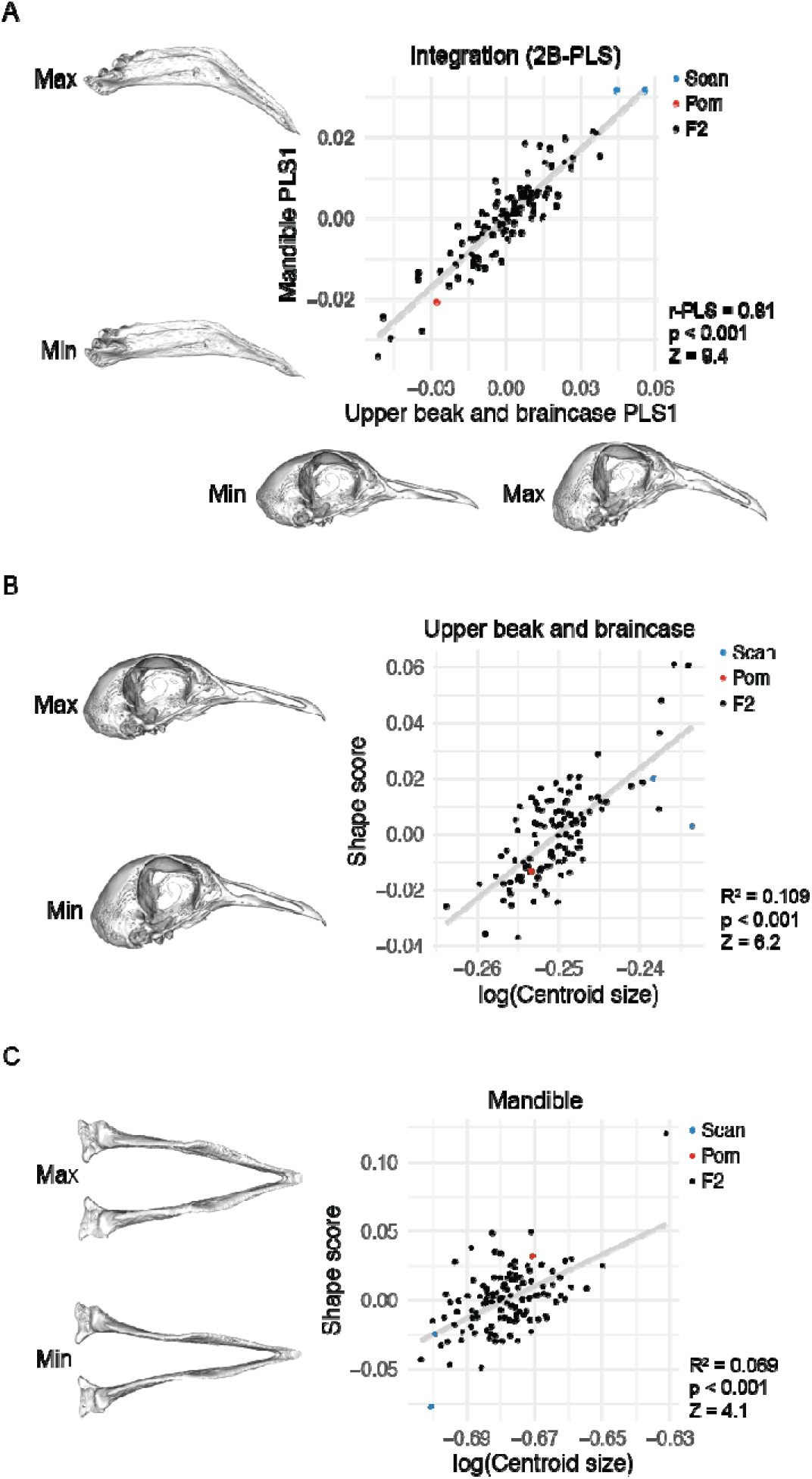
UBB and MAN integration and allometry. (A) UBB PLS1 vs. MAN PLS1 shape. (B) UBB shape ∼ centroid size linear regression. (C) MAN shape ∼ centroid size linear regression. For all panels, minimum and maximum shapes are depicted by warped meshes along corresponding axis. Shape changes were magnified 2x to aid visualization.

**Supplemental Figure 9.**
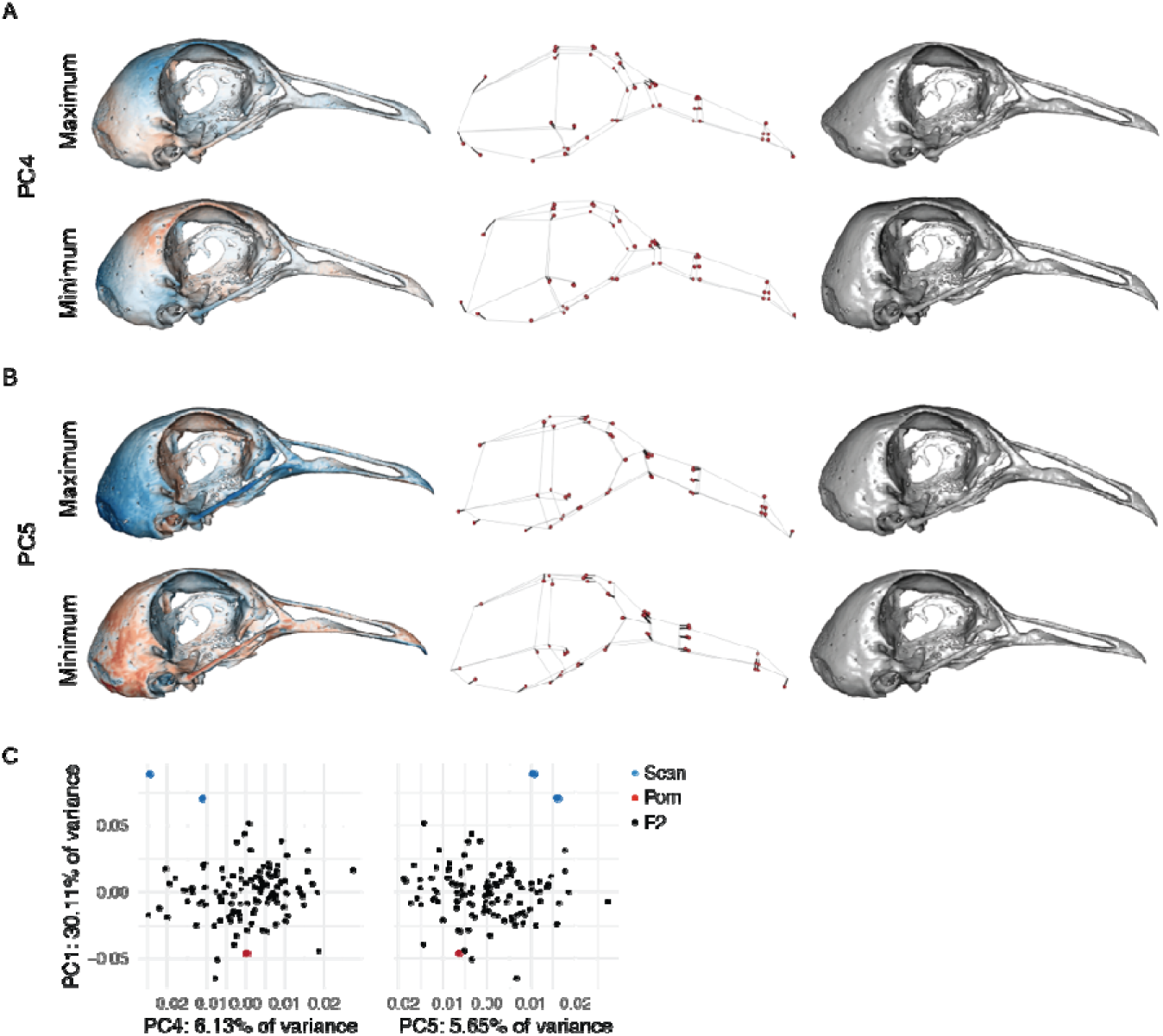
UBB PC4 and PC5 shape variation. (A-B) Minimum and maximum UBB PC4 (A) and PC5 (B) shapes, visualized as heatmaps (left), wireframes (center), and warped meshes (right). For wireframes and meshes, UBB PC4 and PC5 shape is magnified 3x to aid visualization. (C) PCA plots of UBB PC1 vs. PC4 (left) and PC5 (right).

**Supplemental Figure 10.**
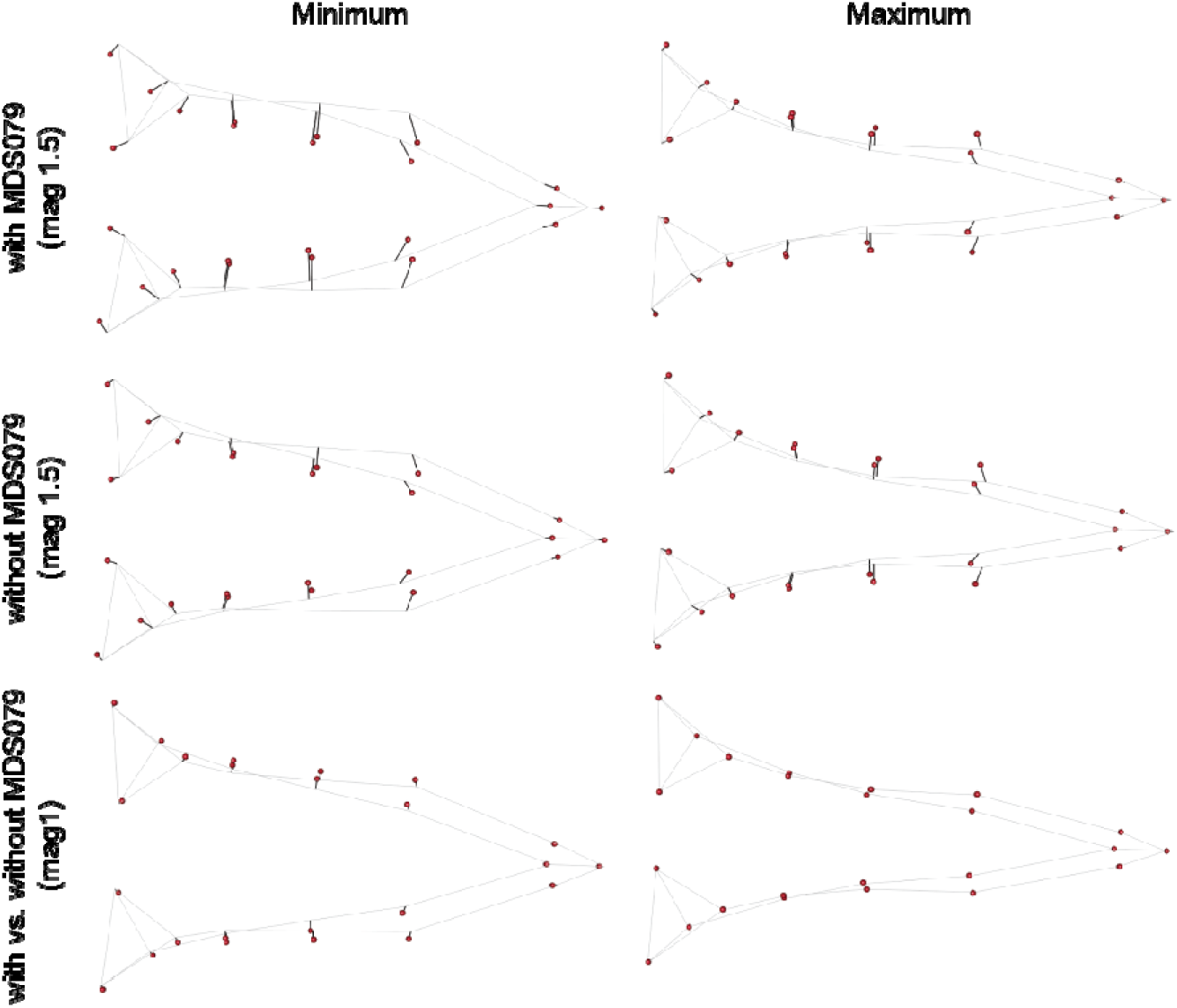
MAN PC1 shape with and without MDS079. Dorsal views of MAN wireframes showing minimum (left) and maximum (right) PC1 shapes if MDS079 is included (top panel) or excluded (center panel) from the geometric morphometric analysis. MDS079 had an exceptionally wide mandible and was an outlier from the rest of the F2 population along the MAN PC1 axis (see PCA plot in Figure 4B). Although inclusion of MDS079 changed the magnitude of the PC1 axis, it had virtually no effect on the shape described by MAN PC1, thus it was kept in all downstream analyses.

**Supplemental Figure 11.**
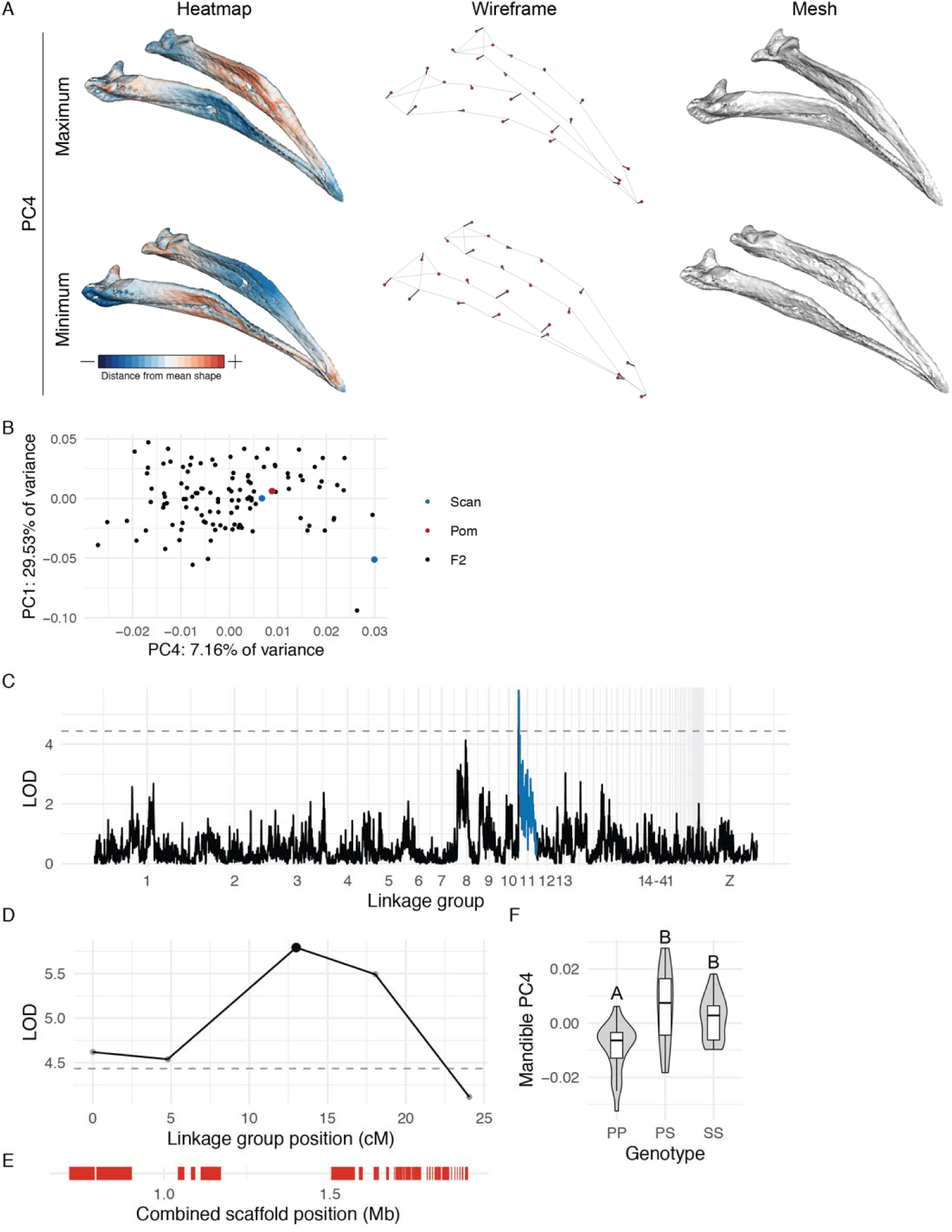
MAN PC4 shape variation and associated QTL. (A) Minimum and maximum MAN PC4 shapes, visualized as heatmaps (left), wireframes (center), and warped meshes (right). For wireframes and meshes, shape is magnified 3x to aid visualization. (B) PCA plots of MAN PC1 vs. PC4. (C) Genome-wide QTL scan for MAN PC4. (D) MAN PC4 LOD support interval for QTL on LG11. (E) Genes in LG11 QTL interval. (F) LG11 QTL effect plot. Letters denote significance groups, p-values determined via Tukey test: PP vs. PS = 2.2e-06, PP vs. SS = 2.2e-03. P = allele from Pom founder, S = allele from Scan founder.

**Supplemental Figure 12.**
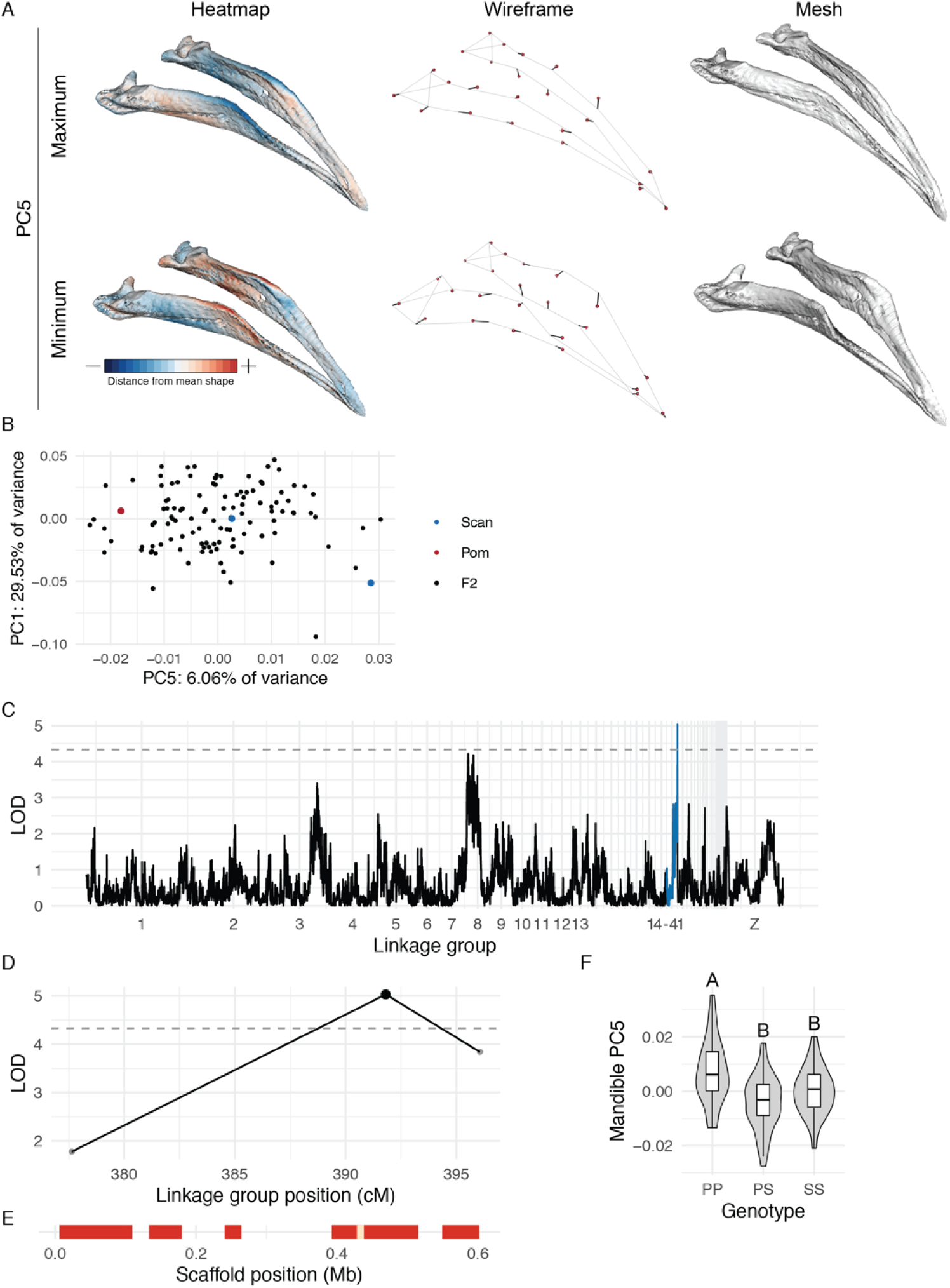
MAN PC5 shape variation and associated QTL. (A) Minimum and maximum MAN PC5 shapes, visualized as heatmaps (left), wireframes (center), and warped meshes (right). For wireframes and meshes, shape is magnified 3x to aid visualization. (B) PCA plots of MAN PC1 vs. PC5. (C) Genome-wide QTL scan for MAN PC5. (D) MAN PC5 LOD support interval for QTL on LG20. (E) Genes in LG20 QTL interval. (F) LG20 QTL effect plot. Letters denote significance groups, p-values determined via Tukey test: PP vs. PS = 1.3e-05, PP vs. SS = 1.9e-02. P = allele from Pom founder, S = allele from Scan founder.

**Supplemental Figure 13.**
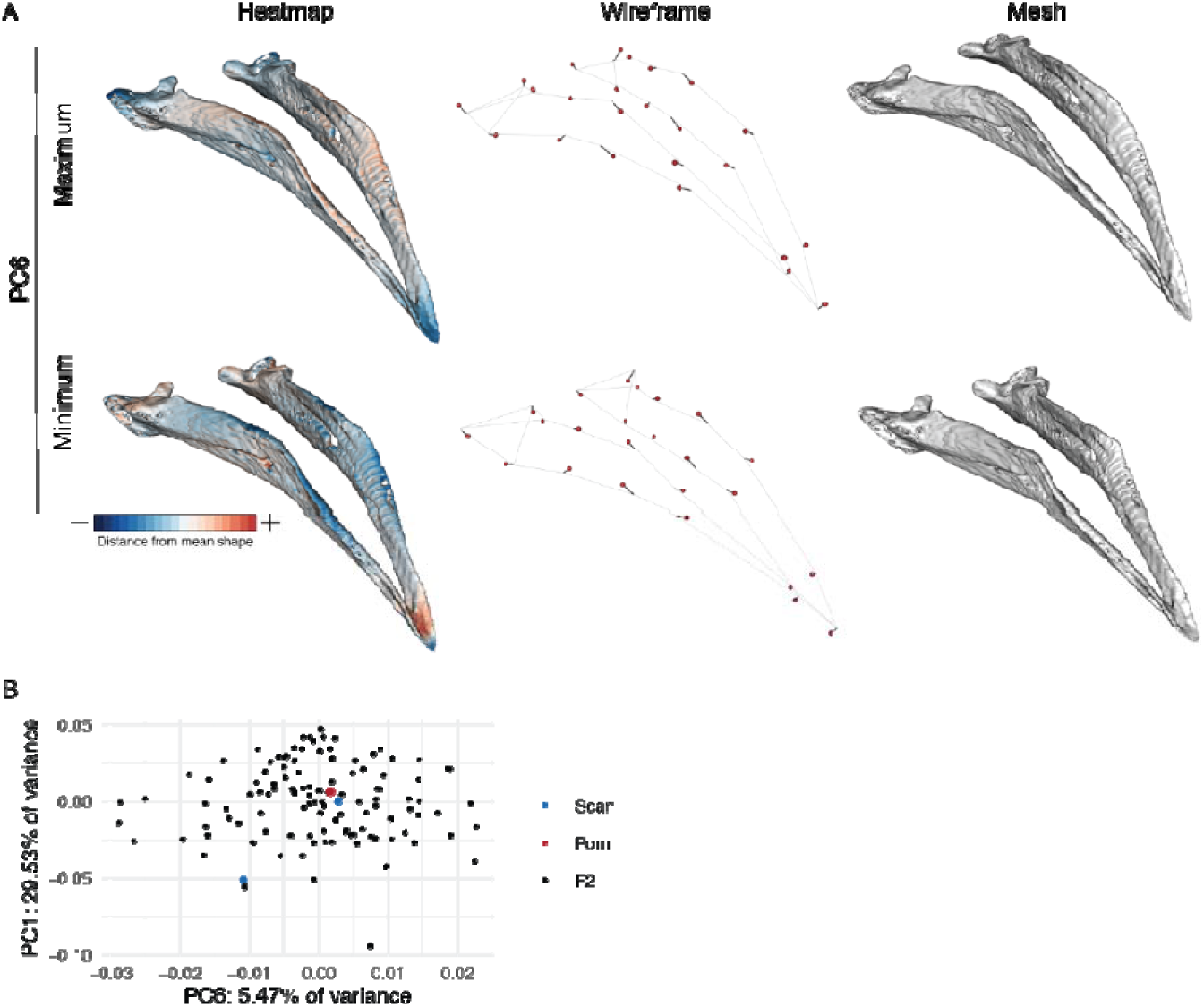
MAN PC6 shape variation. (A) Minimum and maximum MAN PC6 shapes, visualized as heatmaps (left), wireframes (center), and warped meshes (right). For wireframes and meshes, shape is magnified 3x to aid visualization. (B) PCA plots of MAN PC1 vs. PC6.

**Supplemental Figure 14.**
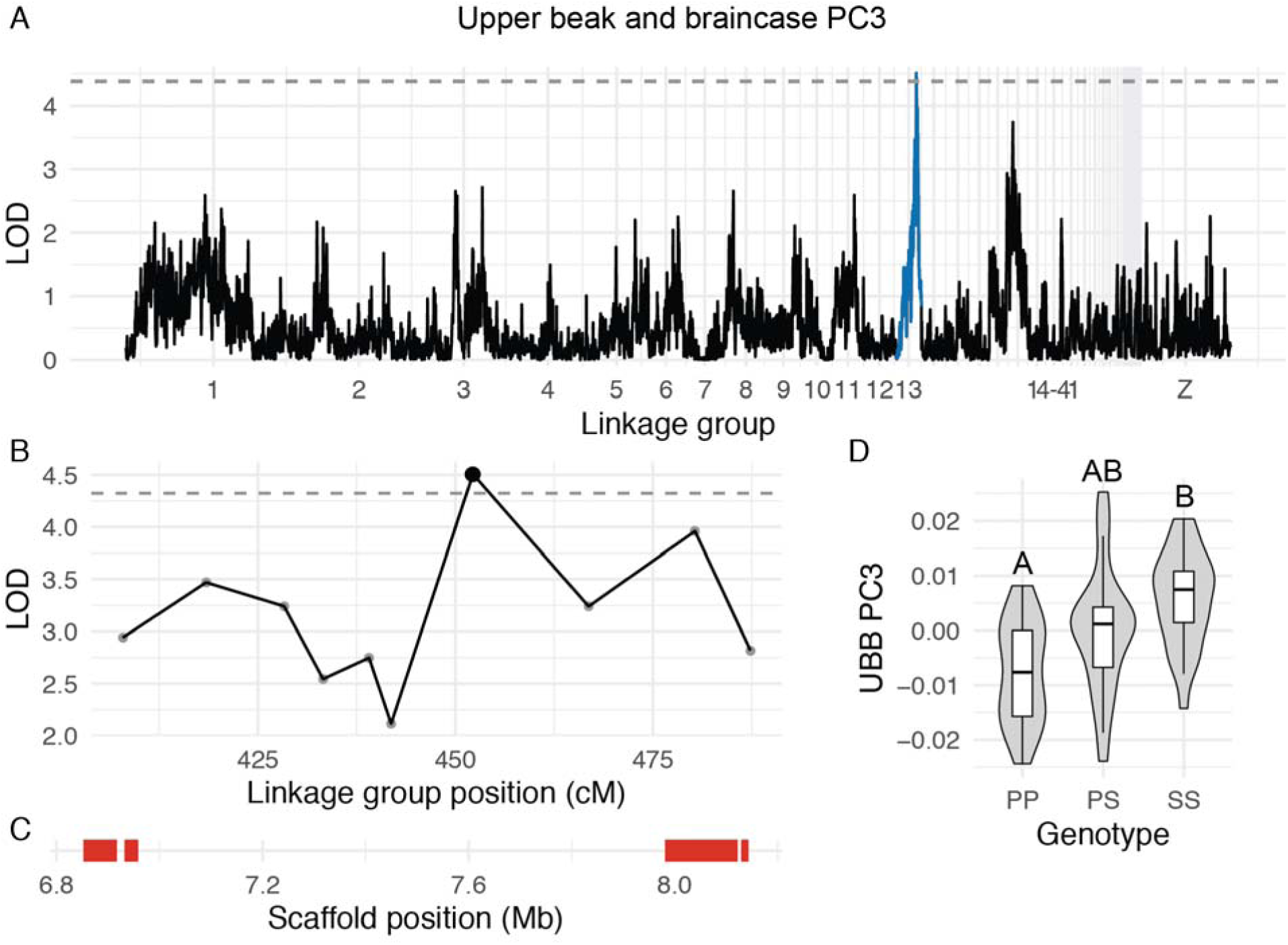
QTL associated with UBB PC3. (A) Genome-wide QTL scan for UBB PC3. (B) UBB PC3 LOD support interval for QTL on LG13. (C) Genes in LG13 QTL interval. (D) LG13 QTL effect plot. Letters denote significance groups, p-values determined via Tukey test: PP vs. SS = 5.3e-04. P = allele from Pom founder, S = allele from Scan founder.

**Supplemental Figure 15.**
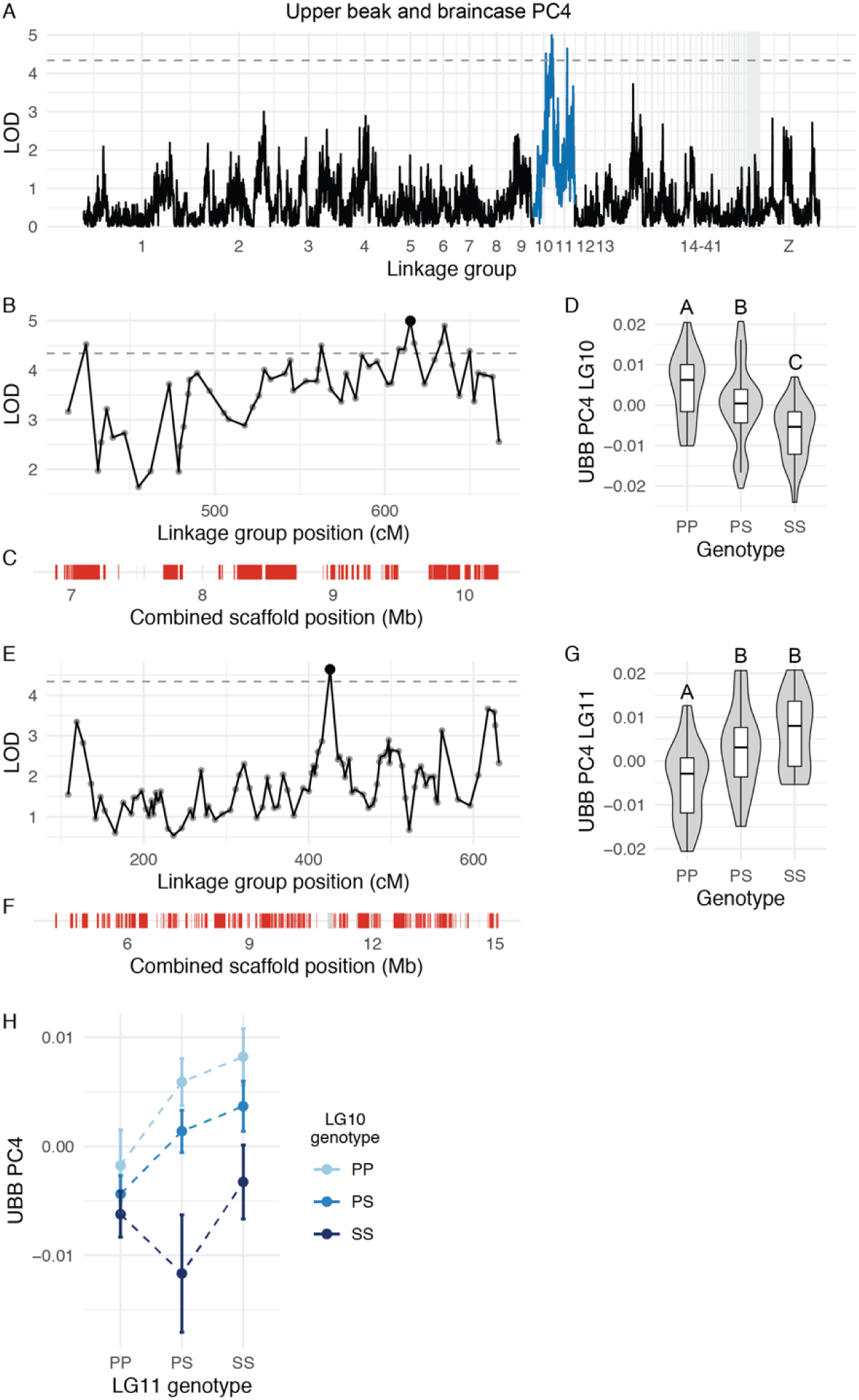
QTL association with UBB PC4. (A) Genome-wide QTL scan for UBB PC4. (B) UBB PC3 LOD support interval for QTL on LG10. (C) Genes in LG10 QTL interval. (D) LG10 QTL effect plot. Letters denote significance groups, p-values determined via Tukey test: PP vs. PS = 1.6e-02, PP vs. SS = 2.4e-05, PS vs. SS = 2.5e-02. (E) LOD support interval for LG11 QTL. (F) Genes in LG11 QTL support interval. (G) LG11 QTL effect plot. Letters denote significance groups, p-values determined via Tukey test: PP vs. PS = 2.4e-03, PP vs. SS = 7.5e-05. (H) Interaction between LG10 and LG11 QTL. P = allele from Pom founder, S = allele from Scan founder.

**Supplemental Movie 1. UBB PC1 shape variation.** Shape change magnified 1.5x.

**Supplemental Movie 2. UBB PC2 shape variation.** Shape change magnified 2x.

**Supplemental Movie 3. UBB PC3 shape variation.** Shape change magnified 3x.

**Supplemental Movie 4. UBB PC4 shape variation.** Shape change magnified 3x.

**Supplemental Movie 5. UBB PC5 shape variation.** Shape change magnified 3x.

**Supplemental Movie 6. MAN PC1 shape variation.** Shape change magnified 1.5x.

**Supplemental Movie 7. MAN PC2 shape variation.** Shape change magnified 1.5x.

**Supplemental Movie 8. MAN PC3 shape variation.** Shape change magnified 2x.

**Supplemental Movie 9. MAN PC4 shape variation.** Shape change magnified 3x.

**Supplemental Movie 10. MAN PC5 shape variation.** Shape change magnified 3x.

**Supplemental Movie 11. MAN PC6 shape variation.** Shape change magnified 3x.

**Supplemental Table 1. Description of skull and jaw landmarks.**

**Supplemental Table 2. Landmark pairs used for skull and jaw linear measurements.**

**Supplemental Table 3. Genes in the beak width and depth LG1 QTL interval.**

**Supplemental Table 4. Genes in the beak depth and mandible width LG8 QTL interval.**

**Supplemental Table 5. Genes in the braincase length LG2 QTL interval.**

**Supplemental Table 6. Genes in the braincase width LG5 QTL interval.**

**Supplemental Table 7. Genes in the mandible length LG10 QTL interval.**

**Supplemental Table 8. Genes in the UBB PC2 LG3 QTL interval.**

**Supplemental Table 9. Genes in the UBB PC3 LG13 QTL interval.**

**Supplemental Table 10. Genes in the UBB PC4 LG10 QTL interval.**

**Supplemental Table 11. Genes in the UBB PC4 LG11 QTL interval.**

**Supplemental Table 12. Genes in the MAN PC3 LG2 QTL interval.**

**Supplemental Table 13. Genes in the MAN PC3 LG3 QTL interval.**

**Supplemental Table 14. Genes in the MAN PC4 LG11 QTL interval.**

**Supplemental Table 15. Genes in the MAN PC5 LG20 QTL interval.**

**Supplemental Table 16. Multi-locus QTL model associated with UBB PC1 shape variation.**

**Supplemental Table 17. Multi-locus QTL model associated with MAN PC1 shape variation.**

